# Costs, Benefits, Parasites and Mutualists: The Use and Abuse of the Mutualism–Parasitism Continuum Concept for *Epichloë* Fungi

**DOI:** 10.1101/2021.04.21.440766

**Authors:** Jonathan A. Newman, Sierra Gillis, Heather A. Hager

## Abstract

The species comprising the fungal endophyte genus *Epichloë* are symbionts of cool season grasses. About half the species in this genus are strictly vertically transmitted, and evolutionary theory suggests that these species must be mutualists. Nevertheless, Faeth and Sullivan (e.g. 2003) have argued that such vertically transmitted endophytes are “usually parasitic,” and Müller and Krauss (2005) have argued that such vertically transmitted endophytes fall along a mutualism–parasitism continuum. These papers (and others) have caused confusion in the field. We used a systematic review to show that close to half of the published papers in this field incorrectly categorize the interaction. Here, we develop the argument that advantages and disadvantages are not the same things as mutualism and parasitism and that experimental evidence must be interpreted in the context of theory. We argue that, *contra* Faeth and Sullivan, it is highly unlikely that such strictly vertically transmitted endophytes can be parasites, and that, *contra* Müller and Krauss, it is inappropriate to apply the continuum concept to strictly vertically transmitted endophytes. We develop a mathematical model to support and illustrate our arguments. We use the model to clarify that parasitism *requires* some degree of horizontal transmission, and that while it is appropriate to use the continuum concept when referring to the genus as a whole, or to species that possess horizontal transmission, we argue that it is inappropriate to use the concept when referring to species that are strictly vertically transmitted.

## 1. Introduction

In this paper, we are concerned with the nature of the ecological interaction between fungal species in the genus *Epichloëe* and their grass host plants. In particular, we consider the existence of a ‘mutualism–parasitism continuum’ for these species. We begin by defining some terms. Within the context of pairwise species interactions, there are ‘consumer interactions’ in which one interacting species benefits and the other is harmed (+, −). These are commonly termed *predation, parasitism, herbivory*, etc. In *commensalism*, one species benefits and the other is not affected (+, 0). In *ammensalism*, one species is harmed and the other is not affected (−, 0). And finally, *mutualism* occurs when both species benefit from the interaction (+, +). We distinguish the use of these terms from fitness *costs* and *benefits*. The costs and benefits of a particular species interaction may shift with species ontogeny, environment, or climate. There may be times during the lifespan of an individual where the costs outweigh the benefits, and other times where the benefits outweigh the costs. As Ewald (1987, p. 295) succinctly put it:

> “The net effect on the host’s inclusive fitness of all harmful and beneficial characteristics is the theoretical criterion used to assign the labels parasitism, commensalism, or mutualism.”

That is, it is the integration of the fitness benefits minus the costs, over the lifespan of the individual, that determines the signs used above to describe the pairwise interaction. Furthermore, we stipulate that in order to *conclusively*, empirically demonstrate the nature of such a pairwise interaction, one would need to measure and compare individual fitness over the lifetime of the individuals involved. Short-term experiments that measure an aspect of, or a surrogate of, fitness provide some information about the immediate costs and benefits of the relationship, but such experiments do not demonstrate the underlying nature of the interaction. Since individual experiments rarely actually measure lifetime fitness, conclusions as to the nature of the interaction are most often *inferences* based on empirical demonstrations of costs and benefits coupled with evolutionary and ecological theory.

### 1.1. Epichloë Fungi

The grass subfamily Poöideae arose 30–40 million years ago, in conjunction with a monophyletic clade of systemic endophytic fungi in the genus *Epichloë*. These fungi colonize most aboveground grass tissues, but are usually absent from root tissues (see Fig. 1). Some *Epichloë* species cause ‘choke disease’ whereby the fungus suppresses the host’s seed production, using the culms as a site to produce fungal ascospores that disperse to infect new host plants (see Fig. 1). However, many *Epichloë* are also able to asymptomatically colonize the grass ovary, ovule and embryo without damaging the seed, and transmit themselves vertically from one host generation to the next. Indeed, some *Epichloë* exhibit choke disease on some tillers, and seed colonization on other tillers. Thus, these species simultaneously undergo sexual recombination combined with horizontal transmission between hosts, and clonal reproduction coupled with vertical transmission from one host generation to the next. Still other species do not reproduce sexually, and transmission to the next generation is *restricted* to vertical transmission (Schardl, 2010).

**Figure 1:**
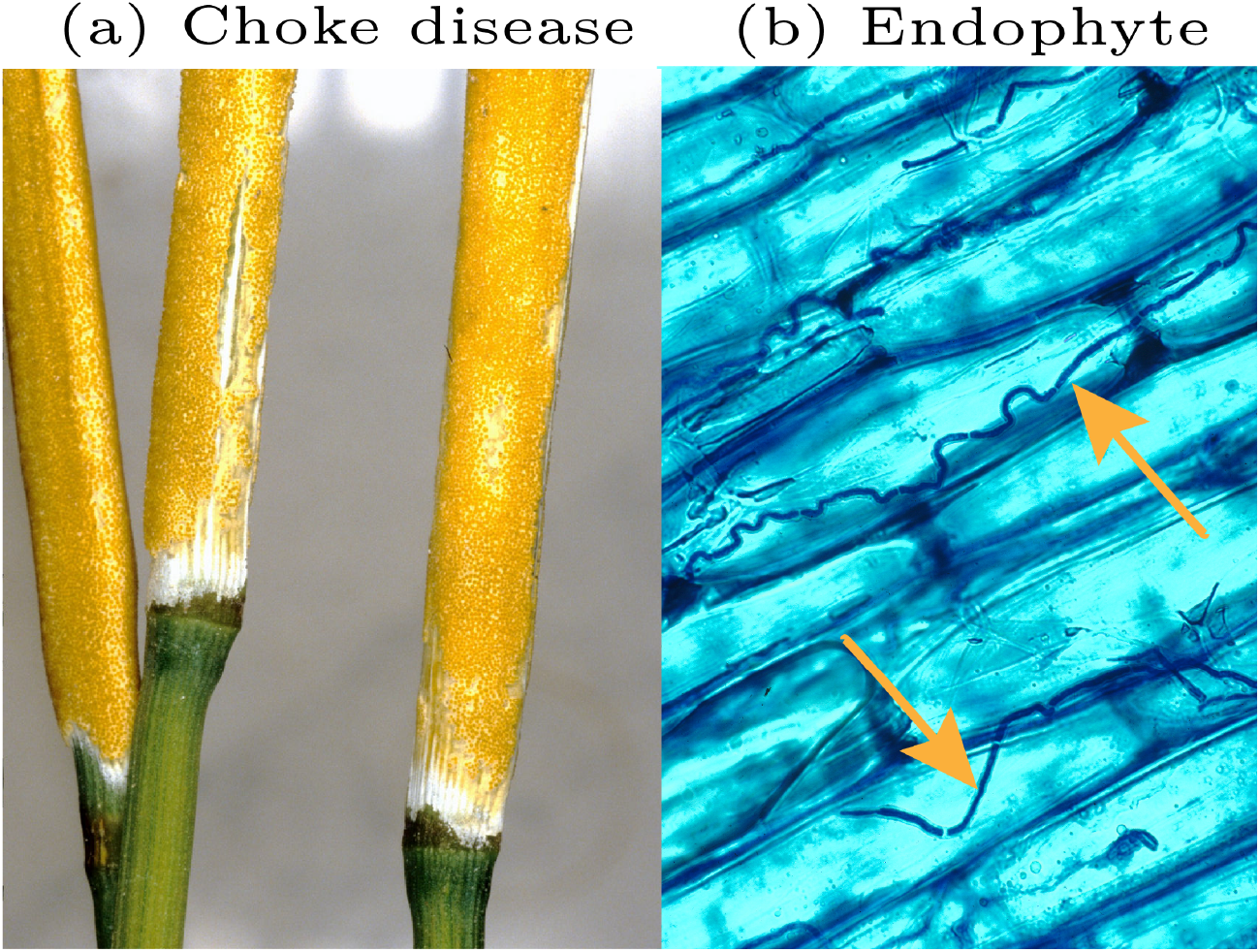
(a) *Epichloë typhina* stromata on grass host, a condition known as ‘choke disease’. Photo copyright 2013 by George Baron; used here with permission. http://hdl.handle.net/10214/5750. (b) Blue-stained micrograph (400 × magnification) of *Epichloë coenophiala* hyphae (orange arrow) inhabiting the intercellular spaces of *Schedonorus arundinaceus* (Schreb.) Dumort. leaf sheath tissue. Photo by Nick Hill, 2006. This image is a work of a United States Department of Agriculture employee, taken or made as part of that person’s official duties. As a work of the U.S. federal government, the image is in the public domain. https://bit.ly/2D7xqKs.

The most recent taxonomic revision of this fungal genus (Leuchtmann et al., 2014) concludes that it comprises 34 species, 3 subspecies, and 6 varieties (i.e. 43 distinct lineages). Whether these species reproduce: only sexually (2 species plus 1 subspecies), only asexually (23 species plus 5 varieties), or both sexually and asexually (9 species plus 2 subspecies) largely determines their mode(s) of transmission into the next generation. Sexual reproduction in these species provides the ability for horizontal transmission of the fungus from one individual host plant to another. Strictly asexually reproducing species seem to be limited exclusively to vertical transmission into the next generation. See Table 1 for details.

**Table 1:** Constructed from information contained in Leuchtmann et al. (2014). Shown are the 34 species, 3 subspecies and 6 varieties of *Epichloë* species, cross classified by whether they are teleomorphic-types or anamorphic-types, and whether they are the result of interspecific hybridization or not. For each taxa, we indicate the presence of sexual reproduction and/or vertical transmission with ✓. The 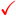 indicates that the reproduction mode occurs on some of the known hosts but not all (e.g., *E. bromicola* transmits itself vertically on 6 of the known host species, but apparently not in the other two). The + indicates that there may be multiple hosts from the same genus or that one of the hosts is known only to genus level. ‘not obs.’ indicates that the transmission mode has not been observed. The bold denotes taxa formerly classified in the genus *Neotyphodium*. See supplementary Table 1 in Leuchtmann et al. (2014) for further details.

|  | Non-Hybrid | Sexual | Vertical | Hosts | Hybrid | Sexual | Vertical | Hosts |
| --- | --- | --- | --- | --- | --- | --- | --- | --- |
| Telomorphic | *E. amarillans* | ✓ | ✓ | 6 | *E. liyangensis* | ✓ | ✓ | 1 |
|  | *E. baconii* | ✓ | absent | 5 |  |  |  |  |
|  | *E. brachyelytri* | ✓ | ✓ | 1 |  |  |  |  |
|  | *E. bromicola* | ✓ | ✓ | 8 |  |  |  |  |
|  | *E. elymi* | ✓ | ✓ | 2+ |  |  |  |  |
|  | *E. festucae* | ✓ | ✓ | 2+ |  |  |  |  |
|  | var *lolii* | not obs. | ✓ | 1 |  |  |  |  |
|  | *E. glyceriae* | ✓ | absent | 1+ |  |  |  |  |
|  | *E. sylvatica* | ✓ | ✓ | 2 |  |  |  |  |
|  | sub *pollinensis* | ✓ | ✓ | 1 |  |  |  |  |
|  | *E. typhina* | ✓ | ✓ | 10 |  |  |  |  |
|  | var *ammophilae* | not obs. | not obs. | 1 |  |  |  |  |
|  | sub *clarkii* | ✓ | absent | 1 |  |  |  |  |
|  | sub *poae* | ✓ | ✓ | 4 |  |  |  |  |
|  | var *aonikenkana* | not obs. | ✓ | 1 |  |  |  |  |
|  | var *canariensis* | not obs. | ✓ | 1 |  |  |  |  |
|  | var *huerfana* | not obs. | ✓ | 1 |  |  |  |  |
| Anamorphic | *E. aotearoae* | not obs. | ✓ | 1 | *E. australiensis* | not obs. | ✓ | 1 |
|  | *E. gansuensis* | not obs. | ✓ | 3 | *E. cabralii* | not obs. | ✓ | 1 |
|  | var *inebrians* | not obs. | ✓ | 1 | *E. canadensis* | not obs. | ✓ | 1 |
|  | *E. mollis* | ✓ | ✓ | 1 | *E. chisosa* | not obs. | ✓ | 1 |
|  | *E. sibirica* | not obs. | ✓ | 1 | *E. coenophiala* | not obs. | ✓ | 1 |
|  | *E. stromatolonga* | not obs. | ✓ | 1 | *E. danica* | not obs. | ✓ | 1 |
|  |  |  |  |  | *E. disjuncta* | not obs. | ✓ | 1 |
|  |  |  |  |  | *E. funkii* | not obs. | ✓ | 1 |
|  |  |  |  |  | *E. guerinii* | not obs. | ✓ | 2 |
|  |  |  |  |  | *E. hordelymi* | not obs. | ✓ | 1 |
|  |  |  |  |  | *E. melicicola* | not obs. | ✓ | 2 |
|  |  |  |  |  | *E. occultans* | not obs. | ✓ | 1+ |
|  |  |  |  |  | *E. pampeana* | not obs. | ✓ | 1 |
|  |  |  |  |  | *E. schardlii* | not obs. | ✓ | 1 |
|  |  |  |  |  | *E. siegelii* | not obs. | ✓ | 1 |
|  |  |  |  |  | *E. sinica* | not obs. | ✓ | 1 |
|  |  |  |  |  | *E. sinofestuae* | not obs. | ✓ | 1 |
|  |  |  |  |  | *E. tembladerae* | not obs. | ✓ | 12 |
|  |  |  |  |  | *E. uncinata* | not obs. | ✓ | 1 |

### 1.2. Parasitism versus mutualism

Sexual reproduction often involves the production of a stromatal stage that ‘chokes’ some of the host plant’s reproductive tillers (see Fig. 1). It is often presumed that these interactions are parasitic because of the obvious fitness loss suffered by the host plant. However, most, if not all of the *Epichloë* lineages produce plant secondary metabolites, alkaloids, that can defend the host plant against some herbivores and pathogens (see e.g. Schardl et al., 2013). So it is unclear what the balance of *lifetime* costs and benefits is in these sexually reproducing lineages. Thus, it is an open question as to whether these lineages are parasites or mutualists, and whether they may be either or both in different environments.

As previously mentioned, the asexual species seem to be restricted to vertical transmission. Endosymbionts that are strictly vertically transmitted *ought* to evolve into mutualists if they are to maintain themselves in the population (see Section 1.3). Indeed, benefits accruing to the host plants were demonstrated ‘early and often’ (for review of the early literature see e.g. Saikkonen et al., 1998), and the interaction was widely asserted to be a mutualism. This view started to be challenged in the mid-1990s when Stan Faeth and his students began studying the native grass *Festuca arizonica* and its asexual endophyte *Neotyphodium starii*.^1^ From early on in their research, Faeth and his colleagues commonly failed to find the widely reported benefits to the host plants of the endophyte infection (see e.g. Lopez et al., 1995) and indeed found costs of harboring the endophyte. The lack of agreement needed an explanation, and one possible reason for the difference that Faeth and colleagues hit upon early on was that they were studying a ‘native’ grass-endophyte pair in natural settings, while much of the early work on the epichloids in general was done on introduced agricultural grasses paired with their non-native *Epichloë* species in managed agroecosystems. Indeed, one can see hints of this explanation forming in the last lines of Faeth’s earliest paper on the subject (Lopez et al., 1995, p. 1579, emphasis added):

> “Our results, however, suggest the effect on herbivores may not be as widespread as currently believed (Clay et al. 1983, Carroll 1988, Clay 1988, Strong 1988), *or the effects are more variable in native grass–herbivore interactions*.”

As Faeth’s body of work began to grow in this field, seemingly so did his conviction that what he and his students were seeing was not the mutualism predicted by evolutionary theory and supported by empirical evidence in the agricultural systems. He began publishing papers with provocative titles such as:

> “*Neotyphodium* in native populations of Arizona Fescue: A nonmutualist?” (Faeth et al., 1997),
>
> “*Neotyphodium* in Arizona fescue: A necessary symbiont?” (Sullivan and Faeth, 1999),
>
> “What maintains high levels of *Neotyphodium* endophytes in native grasses? A dissenting view and alternative hypotheses” (Faeth et al., 2000),
>
> “Fungal endophytes: Common host plant symbionts but uncommon mutualists” (Faeth and Fagan, 2002),
>
> “Are endophytic fungi defensive plant mutualists?” (Faeth, 2002), culminating in the particularly assertive:
>
> “Mutualistic asexual endophytes in a native grass are usually parasitic” (Faeth and Sullivan, 2003).

He continued to develop these ideas throughout these papers, for example (Faeth, 2002, p. 25):

> “Endophytic fungi, especially asexual, systemic endophytes in grasses, are generally viewed as plant mutualists, mainly through the action of mycotoxins, such as alkaloids in infected grasses, which protect the host plant from herbivores. Most of the evidence for the defensive mutualism concept is derived from studies of agronomic grass cultivars, which may be atypical of many endophyte–host interactions. I argue that endophytes in native plants, even asexual, seed-borne ones, rarely act as defensive mutualists. In contrast to domesticated grasses where infection frequencies of highly toxic plants often approach 100%, natural grass populations are usually mosaics of uninfected and infected plants. The latter, however, usually vary enormously in alkaloid levels, from none to levels that may affect herbivores. This variation may result from diverse endophyte and host genotypic combinations that are maintained by changing selective pressures, such as competition, herbivory and abiotic factors. Other processes, such as spatial structuring of host populations and endophytes that act as reproductive parasites of their hosts, may maintain infection levels of seed-borne endophytes in natural populations, without the endophyte acting as a mutualist.”

As Faeth has continued to add to this argument through the years, others too started detecting costs of the endophyte to the host grass (e.g. Cheplick et al., 2000, Cheplick, 2007, 2008, Härri et al., 2008a, b, Fuchs et al., 2013). Taken together, Faeth’s arguments and the publication of other papers demonstrating costs have given rise to an (albeit minority) view that the strictly asexual vertically transmitted *Epichloë*-grass interactions form a ‘mutualism–parasitism continuum’ (Müller and Krauss, 2005). In a systematic survey of the literature (323 papers, see Appendix A), we found that for strictly vertically transmitted *Epichloë* species, 58% of the papers considered the interaction to be a mutualism, but 30% considered it at least possible that the interaction could indeed be a parasitism (see Table 2 for details). And this view persists today (see e.g. Raman and Suryanarayanan, 2017, and Fig. A.1).

**Table 2:** Summary of a systematic review of 323 papers (see Appendix A for details). We classified papers by the relationship claimed by the authors and cross-classified these claims by the mode of transmission for the *Epichloë* species being studied in each case (see Table 1. HT: horizontal transmission, VT: vertical transmission. *Continuum* denotes a claim by the authors that the *Epichloë* species and host plant interaction is best described as belonging on a continuum between parasitic and mutualist. *Admit the possibility* denotes a claim by the author that it is possible that the species interaction could be parasitic. *Not stated* denotes that the authors made no statement at all as to the nature of the interspecific interaction. The cells with red text and and ‘*’ represent classification errors. As we show elsewhere in this paper, it is incorrect to classify species that possess horizontal transmission as strictly mutualist (they may be, they also may be parasites). Conversely, it is incorrect to classify strictly vertically transmitted species as anything other than mutualists. See Discussion and Appendix A for more details.

| Relationship Claimed | Transmission mode |  |
| --- | --- | --- |
|  | HT or HT+VT | VT only |
| Parasitic | 3 | 2* |
| Continuum | 14 | 40* |
| Admit the possibility | 6 | 20* |
| Mutualism | 80* | 121 |
| Not stated | 12 | 25 |

### 1.3. Why is this view controversial?

Cases of a parasite exhibiting only vertical transmission are interesting from an evolutionary perspective. To see why, let: *ρ* be the proportional change in host fecundity relative to an uninfected host; *ψ* be the proportional change in the probability of survival from seed through to reproduction; *γ* be the probability an infected mother will infect an average seed; and *δ* be the same probability for the father. Then, from the Fundamental Vertical Transmission Equation, it can be shown that a sufficient condition for the persistence of a parasite exhibiting only vertical transmission is *ρψ*(*γ*+*δ*) > 1, and that outside of the possibly rare condition that *γ* + *δ* > 1.65, it is also a necessary condition (Fine, 1975). By definition, a parasite reduces the lifetime fitness of its host relative to an uninfected conspecific, so *ρψ* < 1. The condition for a vertically transmitted parasite to maintain itself in a population thus requires a sufficiently high transmission rate (*γ* + *δ*) to counterbalance the fitness detriment it causes to its host. In the case of the asexual *Epichloë* species, the fungus has never been shown to be transmitted via the pollen, and therefore *δ* = 0. Even assuming that maternal vertical transmission were perfect (i.e. *γ* = 1) it is not possible for the fungus to maintain itself in the population while still causing a reduction in the lifetime fitness of the host species. That is, the fungus cannot continue to be a parasite, and evolutionary theory thus strongly suggests that such species must evolve into mutualists (Ewald, 1987).

In the face of such theory, those who would argue that these strictly vertically transmitted fungi are not mutualists must provide a mechanism that explains how the fungus is maintained within a plant population, in the face of lifetime fitness costs to the plant. Faeth and Sullivan (2003) offer four possible explanations for their claim that “mutualistic asexual endophytes in a native grass are usually parasitic” (see also Faeth, 2010). We briefly summarize these here. See Faeth and Sullivan for a more detailed presentation.

1. *Mutualistic outcomes are rare but important*. This explanation is an acknowledgement that it is lifetime fitness that matters, not a component of fitness (e.g. seed production), measured over a period of a few months or years. These cool-season perennial grasses can live for tens of years. What happens during a single season, or indeed multiple seasons, matters but has to be viewed in the context of the entire lifetime. This is particularly true if the mutualist advantage is very large in these ‘rare’ years. This defense amounts to an admission that the interaction is indeed a mutualism, when viewed over the whole lifetime of the host plants. We agree that this is a plausible explanation of experimental demonstrations of endophyte induced fitness reductions to the host plant. However, we feel it is misleading to refer to such a situation as a ‘parasitism’ when it is plainly a mutualism. In our view, it is better to refer to such observations as ‘costs.’
2. *Interactions vary with spatial structuring*. This defense imagines a landscape containing multiple local populations of the grass hosts. In these populations, some individuals will be infected with the endophyte, and some individuals will be endophyte-free. Furthermore, it is imagined, there will be random ‘catastrophes’ that temporarily wipe-out some populations, and these now ‘empty’ patches become available for recolonization. If these local populations were closed (i.e. no dispersal between them), *and the endophyte were parasitic*, the whole metapopulation would eventually lose the endophyte. However, if dispersal between populations is sufficiently high, and catastrophes are sufficiently common, then it is (mathematically) possible for a parasitic strain of the endophyte to persist in the meta-population (Saikkonen et al., 2002). Intuitively, this works because host plants infected with a parasitic strain of the endophyte would be able to disperse and recolonize empty patches (wiped out by the catastrophes) before they are out-competed by endophyte-free plants. This meta-population explanation requires significant dispersal of *seeds* (not pollen) between populations. With some notable exceptions, grass seeds are not known for their long-distance dispersal, typically on the order of tens of centimeters (e.g., Rabinowitz and Rapp, 1981), rather than the longer distances that might be necessary for persistence in a landscape such as is imagined here. Therefore, we do not think that this mechanism seems very plausible, but admittedly the hypothesis awaits experimental evidence one way or the other.
3. *Asexual endophytes are transmitted horizontally*. This hypothesis does not explain how *strictly* vertically transmitted endophytes can be parasites and still persist in the population. Instead, it changes the question. If these asexual fungal endophytes occasionally do transmit themselves horizontally, this may account for how an endophyte might be parasitic and still maintain itself in the population. This explanation seems unlikely since horizontal transmission in these species has not been observed, and, as Faeth and Sullivan admit, the horizontal transmission rate would have to be high enough to overcome the endophyte loss rate, which can be high (Afkhami and Rudgers, 2008, Rudgers et al., 2009). To these caveats, we would add that basic epidemiological theory requires the dispersal rate to be ‘high enough’ to overcome the fitness detriment caused to the host plant (i.e. the parasite must be able to disperse faster than it is killing its host; see e.g. Anderson and May, 1992, Chapter 4). See also Faeth et al. (2007) for a theoretical treatment of this idea and other variations on the theme.
4. *Asexual endophytes retain control over host plant reproduction*. This explanation is an attempt to extend a hypothesis about *Wolbachia* bacteria to the *Epichloë* context. *Wolbachia* form associations with many arthropods and filarial nematodes. In the nematodes, *Wolbachia* form stable mutualistic symbioses. In insects, *Wolbachia* are predominantly vertically transmitted, and are supposedly an evolutionary conundrum, because they seem to violate the theory, described earlier, that heritable symbionts must be mutualists. *Wolbachia* seem to be reproductive parasites, manipulating the reproduction of their arthropod hosts to their own advantage, and in the process often substantially decreasing the host’s fitness. *Wolbachia* have been shown to use cytoplasmic incompatibility, killing or feminization of genetic males, and induction of thelytokous parthenogenesis. The supposed advantage of these manipulations for *Wolbachia* is that they are inherited solely through the female germline (as are vertically transmitted *Epichloë*), and their manipulations increase the proportion of females in the host population, allowing *Wolbachia* to spread without being a mutualist (Zug and Hammerstein, 2015).

In a recent review article, Zug and Hammerstein (2015, p. 92; references omitted) offer this response to the supposed ‘conundrum’:

> “Several arguments can be raised to reconcile the view that vertical transmission leads to mutualistic interactions with the existence of SGEs [selfish genetic elements] such as *Wolbachia*. Obviously, *Wolbachia* can indeed evolve into mutualists, as discussed herein. In the bedbug, for example, *Wolbachia* reside in a bacteriome and supply the host with B vitamins. Secondly, there is broad phylogenetic evidence for recurrent horizontal transmission of *Wolbachia* on evolutionary timescales. Horizontal transmission is likely to be a major reason why *Wolbachia* have not evolved more frequently to mutualists in arthropods. Lastly, it has been argued that SGEs are consistent with the conventional hypothesis if symbiont transmission is measured from the perspective of host genes instead of host organisms. In this gene-centered view of symbiont transmission, host sexual reproduction can be regarded as horizontal transmission of SGEs which allows them to become virulent.”

Turelli (1994) points out that, despite there being no *need* for *Wolbachia* to be a mutualist, there is still selection pressure acting to promote mutualism. A mutualist strain of *Wolbachia* that, in addition to manipulating its host’s reproduction, also conferred a fitness advantage to the host, would spread in a population of *Wolbachia* where such a trait was absent. Zug and Hammerstein (2015, p. 90) point out that “recent years have seen a growing body of evidence suggesting that *Wolbachia* can have positive effects on the fitness of arthropod hosts and thus behave as mutualists.” While the *Wolbachia* conundrum is still being debated, there are several elements of this story that do not seem to apply in the *Epichloë* case. The occasional horizontal transmission in *Wolbachia*, and the lack of evidence for manipulation of the grass host sex ratios are two obvious ones.

### 1.4. The ‘mutualism–parasitism continuum’

Ewald (1987) probably introduced the term *mutualism–parasitism continuum*, but Johnson et al. seem to have popularized the term in their con-sideration of mycorrhizal fungi-host plant interactions.^2^ They described it thus (Johnson et al., 1997, p. 583):

> “Mutualism and parasitism are extremes of a dynamic continuum of species interactions. Mycorrhizal associations are generally at the mutualistic end of the continuum, but they can be parasitic when the stage of plant development or environmental conditions make costs greater than benefits, or possibly when the genotypes of the symbionts do not form ‘win-win’ associations. In natural systems, plant genotypes exist because they successfully propagate more plants, and fungal genotypes exist because they successfully propagate more fungi. Usually, by living together, plants and mycorrhizal fungi improve each other’s probability for survival and reproductive success. But sometimes, and being anthropomorphic, plant ‘interests’ are in conflict with those of the fungi. In the words of Dawkins (1978) “… we must expect lies and deceit, and selfish exploitation of communication to arise whenever the interests of the genes of different individuals diverge.”
>
> The ‘interests’ of plants and mycorrhizal fungi are likely to di-verge in highly managed agricultural systems, where fertilization eliminates shortages of soil nutrients, and plant genotypes are selected by humans and not by millennia of natural selection.”

Although the argument about where on this supposed continuum these epichloid species sit had been going on for quite some time (for a sampling of this literature, see: Clay, 1988, 1991, Saikkonen et al., 1999, Faeth et al., 1997, Faeth and Fagan, 2002, Faeth et al., 2000, 2004, Faeth and Hamilton, 2006, Rudgers et al., 2010, 2012), Müller and Krauss (2005) seem to have introduced the use of the term to the study of epichloids with their paper titled: “Symbiosis between grasses and asexual fungal endophytes”:

> “The symbiosis between vertically transmitted asexual endophytic fungi and grasses is common and generally considered to be mutualistic. Recent studies have accumulated evidence of negative effects of endophytes on plant fitness, prompting a debate on the true nature of the symbiosis. Genetic factors in each of the two partners show high variability and have a range of effects (from positive to negative) on plant fitness. In addition, interacting environmental factors might modify the nature of the symbiosis. Finally, competition and multitrophic interactions among grass consumers are influenced by endophytes, and the effects of plant neighbours or consumers could feedback to affect plant fitness. *We propose a mutualism–parasitism continuum for the symbiosis between asexual endophytes and grasses, which is similar to the associations between plants and mycorrhizal fungi.”* (p. 450; emphasis added)

While debate over the usefulness of the concept continues in the mycorrhizal literature (see e.g., Smith and Smith, 2013), in this paper we argue that the concept is useful to the *Epichloë* situation *only* when discussing the genus as a whole, or when discussing individual species that possess horizontal transmission. We argue that, *contra* Muäller and Krauss, the term is not appropriate for use with strictly vertically transmitted *Epichloë*. There are important biological differences between mycorrhizal fungi and *Epichloë* fungi that make this continuum concept less universal than it is perhaps in the mycorrhizal situation. As Ewald (1987) pointed out, transmission mode is probably the most important difference. In the next section we develop a simple model of each transmission strategy and use this model to draw conclusions about the nature of these *Epichloë*-grass interactions.

## 2. A Model of Endophyte Transmission

In this section we develop a simple continuous-time, ordinary non-linear differential equation model that covers each of the three cases of transmission: horizontal transmission only, vertical transmission only, or both horizontal and vertical transmission. Before we develop the model, we first take care of some housekeeping. The notation used in the model is summarized in Table 3.

**Table 3:**
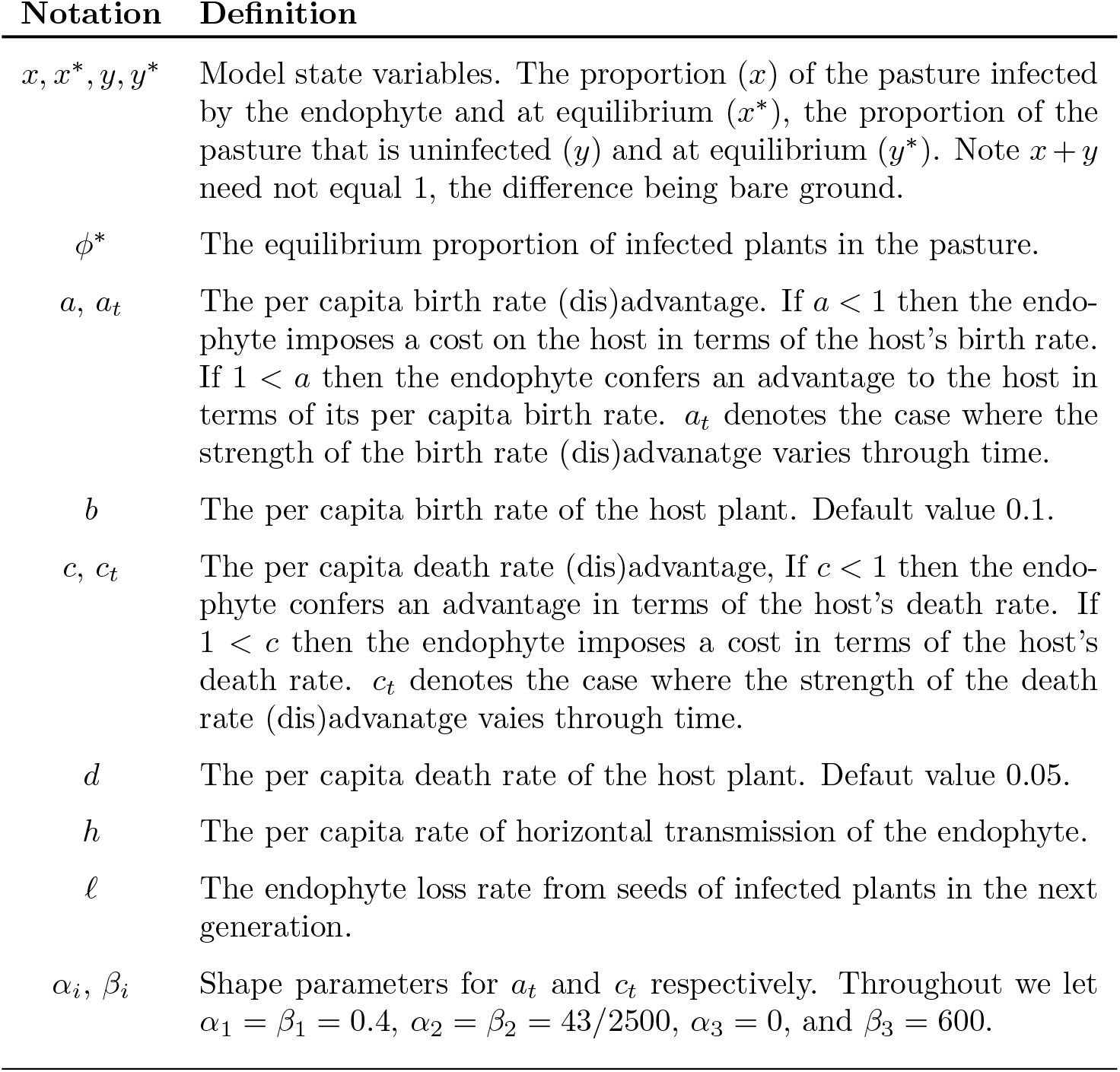
Model definitions. These parameters and state values are used in Eq. (9).

| Notation | Definition |
| --- | --- |
| $x, x^*, y, y^*$ | Model state variables. The proportion ($x$) of the pasture infected by the endophyte and at equilibrium ($x^*$), the proportion of the pasture that is uninfected ($y$) and at equilibrium ($y^*$). Note $x + y$ need not equal 1, the difference being bare ground. |
| $\phi^*$ | The equilibrium proportion of infected plants in the pasture. |
| $a, a_t$ | The per capita birth rate (dis)advantage. If $a < 1$ then the endophyte imposes a cost on the host in terms of the host's birth rate. If $1 < a$ then the endophyte confers an advantage to the host in terms of its per capita birth rate. $a_t$ denotes the case where the strength of the birth rate (dis)advantage varies through time. |
| $b$ | The per capita birth rate of the host plant. Default value 0.1. |
| $c, c_t$ | The per capita death rate (dis)advantage. If $c < 1$ then the endophyte confers an advantage in terms of the host's death rate. If $1 < c$ then the endophyte imposes a cost in terms of the host's death rate. $c_t$ denotes the case where the strength of the death rate (dis)advantage varies through time. |
| $d$ | The per capita death rate of the host plant. Default value 0.05. |
| $h$ | The per capita rate of horizontal transmission of the endophyte. |
| $\ell$ | The endophyte loss rate from seeds of infected plants in the next generation. |
| $\alpha_i, \beta_i$ | Shape parameters for $a_t$ and $c_t$ respectively. Throughout we let $\alpha_1 = \beta_1 = 0.4$, $\alpha_2 = \beta_2 = 43/2500$, $\alpha_3 = 0$, and $\beta_3 = 600$. |

### 2.1. State variables

Let 0 ≤ *x* ≤ 1 be the proportion of a pasture that is infected with the endophyte. Let 0 ≤ *y* ≤ 1 be the proportion of the pasture that is uninfected. At equalibrium, we denote these as *x*\* and *y*\*. We can then define

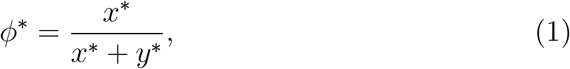

as the proportion of the plants in the pasture that are infected with the endophyte at any equilibrium. Note that *x*+*y* need not equal 1; the difference simply represents bare ground.

### 2.2. Density-dependence

We assume that both birth and death rates are density dependent (e.g. Bullock et al., 1994). Let *b* be the density-*independent* per capita birth rate, and *d* be the density-*independent* death rate. Note that the model we will develop lacks a stage-structure. Here, ‘birth’ means the rate at which individual plants successfully recruit from seeds (i.e. it includes seed and seedling mortality), while ‘death’ means the death of mature plants. We then model the density-*dependent per capita* birth and death rates, respectively, as:

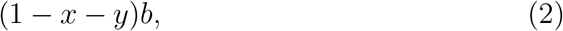

and

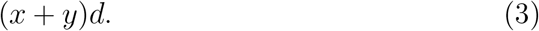

### 2.3. Endophyte (dis)advantage

Throughout the remainder of this paper we refer to the effects of the presence of the endophyte as either an *advantage* or a *disadvantage* to the host plant in terms of the effects the endophyte has, directly or indirectly, on the host’s birth rate and death rate. It would also be appropriate to refer to these as benefits and costs of harbouring the endophyte. We use the term *(dis)advantage* to refer to the effect of the endophyte on the host generally, to denote circumstances in which the these effects might be positive or negative.

Let *a* be the per capita birth rate (dis)advantage. We then model the endophyte infected proportion of the pasture’s per capita birth rate as:

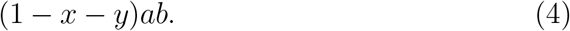

If *a* = 1 then there is neither an advantage nor a disadvantage to the host plant of harboring the endophyte, in terms of the plant’s birth rate. If *a* > 1 then the endophyte is an advantage to the plant (at least in terms of its birth rate) because it increases the birth rate. For example, if *a* =1.2 then the per capita birth rate of infected host plants will be 20% greater than the per capita birth rate of uninfected host plants. And if *a* < 1 then the endophyte imposes a disadvantage on the host plant, at least in terms of its birth rate.

Similarly, let *c* be the per capita death rate (dis)advantage. We model the endophyte infected proportion of the pasture’s per capita death rate as:

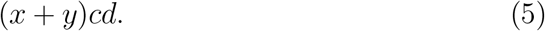

If *c* = 1 the endophyte imposes neither an advantage nor a disadvantage to the host plant. If *c* < 1 then the endophyte is an advantage to the host plant, at least in terms of its per capita death rate, and if *c* > 1 then the endophyte confers a disadvantage to the host plant in terms of death rates. These relationships are summarized in Fig. 2.

**Figure 2:**
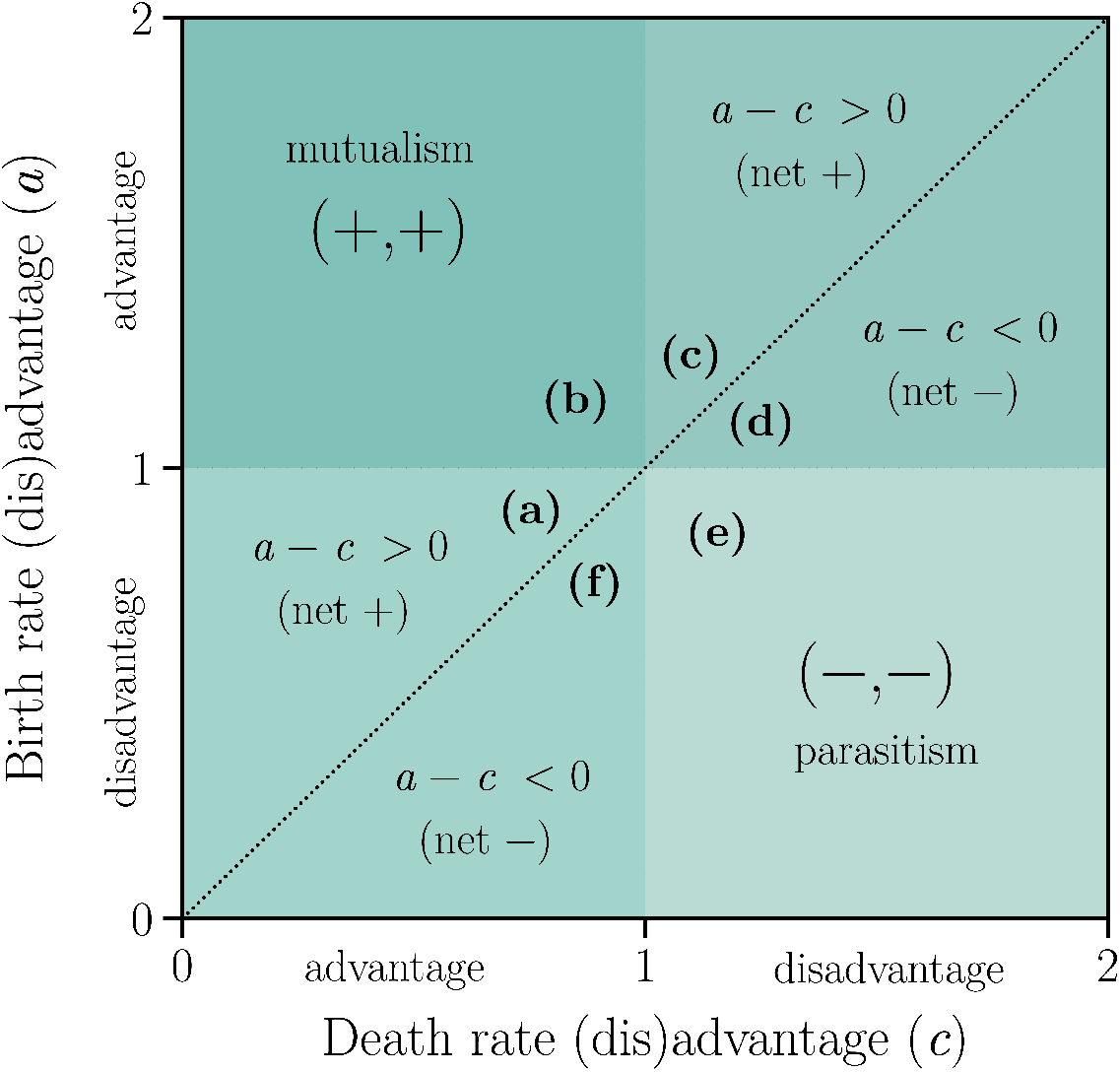
Parameter space. *a* denotes the birth rate (dis)advantage. If 1 < *a* the endophyte confers an advantage to the host plant in terms of the plant’s birth rate. *c* denotes the death rate (dis)advantage. If *c* < 1 the endophyte confers an advantage in terms of the plant’s death rate. The species interaction would certainly be called a *mutualism* in the upper left hand quadrant of the parameter space since the plant enjoys an advantage of the endophyte infection in terms of both its birth and death rates. In the lower right hand quadrant, the species interaction would certainly be called a *parasitism* since the plant suffers a disadvantage of the endophyte infection in terms of both its birth and death rates. The remaining two quadrants represent trade-offs between
an advantage and a disadvantage. In the upper right the endophyte confers an advantage in terms of the plant’s birth rate, but a disadvantage in terms of the plant’s death rate. Whether or not this interaction is a mutualism or a parasitism depends on the relative strengths of the two (i.e. whether or not it falls above the *a* = *c* line or below it, respectively). And similarly for the lower left quadrant except that now the endophyte confers a disadvantage in terms of the plant’s birth rate but an advantage in terms of its death rate. The (a), (b),…(f) denote particular examples that we illustrate in Fig. 3 and elsewhere in the results.

#### 2.3.1. Temporally varying (dis)advantages

Up to now, we have been assuming that *a* and *c* are constants. In reality, endophyte effects vary through time (e.g., Faeth et al., 2006) and interact with the growing environment (e.g., Helander et al., 2016). We think much of the confusion around the mutualism–parasitism continuum derives from such situations. We therefore develop this situation as well. Let

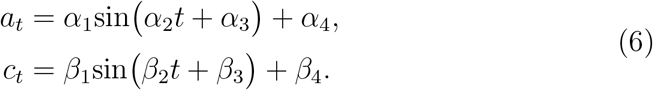

The shape parameters work as follows, *a_t_* oscillates around *α*_4_ ± *α*_1_ and *c_t_* oscillates around *β*_4_ ± *β*_1_. The period of the oscillations is 2*π*/*α*_2_ and 2*π*/*β*_2_. The phase shifts are given by *α*_3_ and *β*_3_.^3^ Note that throughout, for simplicity, we let *α*_3_ = 0, *β*_3_ = 600, and *α*_2_ = *β*_2_ = 43/2500 which yields a period of 365.3 days.

If we integrate Equation 6 over the interval [0, 2*π*/*α*_2_] we get:

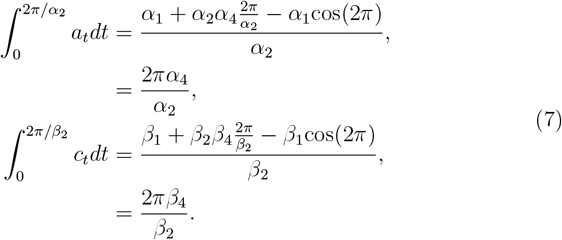

From Eq. (7) we can show that the mean values of *a_t_* and *c_t_* are:

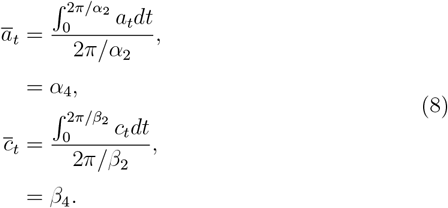

The situation described above is illustrated in Fig. 3. How should we describe the relationships depicted in this figure? There are times of the year when one, the other, or both of the birth and death rates are disadvantaged by the presence of the endophyte. Does this make them parasites? We argue that in Fig. 3a, 3b and 3c the endophytes are mutualists because 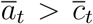, while in Fig. 3d, 3e and 3f the endophytes are parasites because 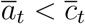 (see also Fig. 2). We provide support for this argument as we develop the model below. We will return to this question in the Discussion section.

**Figure 3:**
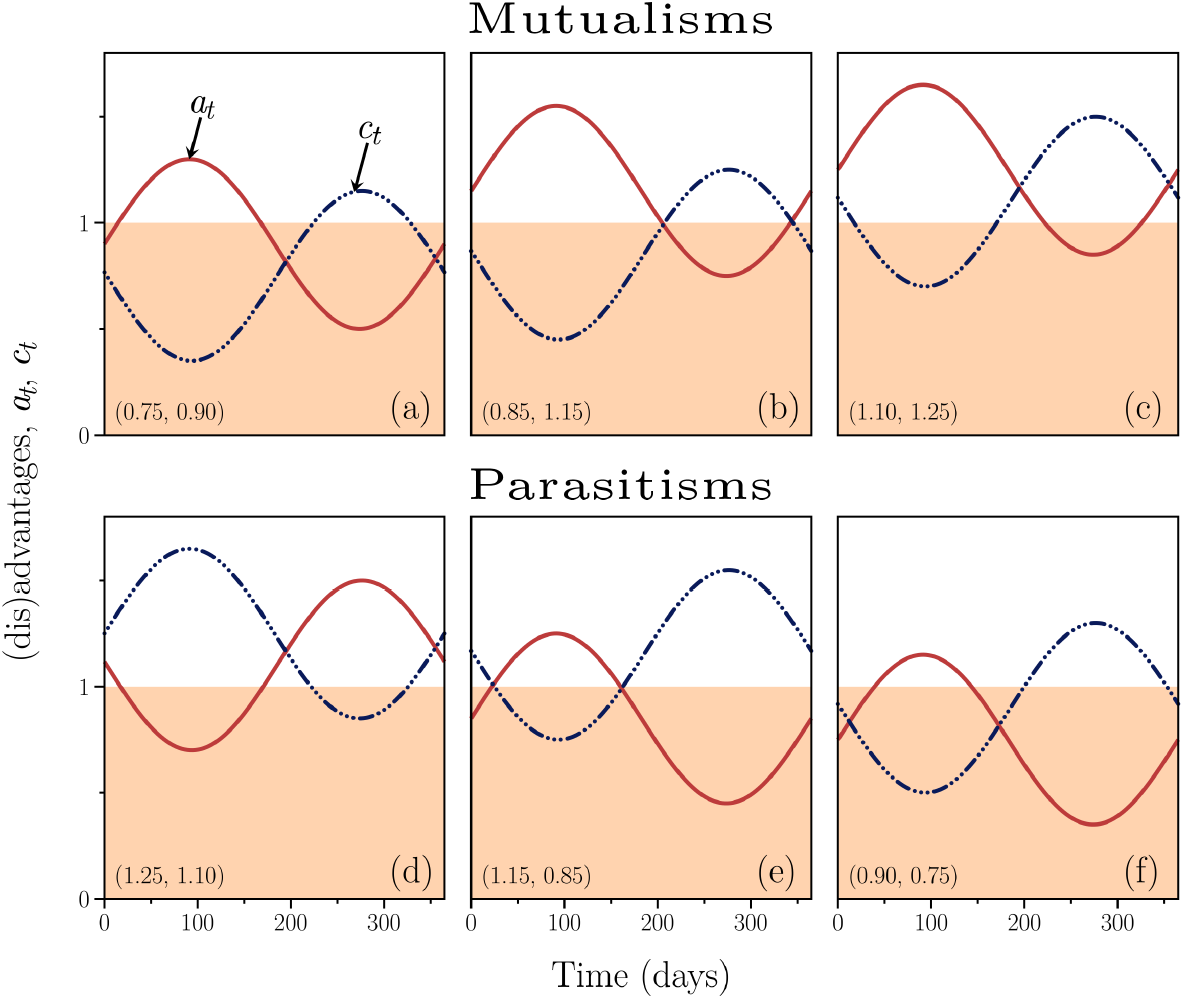
Temporally varying birth and death rate (dis)advantages. The endophyte confers a birth rate advantage when *a_t_* > 1 and a disadvantage otherwise. The endophyte confers a death rate advantage when *c_t_* < 1 and a disadvantage otherwise. In each panel the (#,#) denote the (*β*_4_, *α*_4_) used to generate the figure. In all panels there are times of the year when the endophyte confers a disadvantage in terms of the host’s birth rate and death rate. In (a), (b) and (c) the advantages of infection outweigh the disadvantages averaged over the year. In (d), (e), and (f) the disadvantages outweigh the advantages averaged over the year.

### 2.4. Endophyte loss

Let 0 ≤ *ℓ* ≤ 1 be the per capita loss rate of the endophyte, and conversely, 1–*ℓ* is the transmission efficiency rate. Endophyte loss occurs through several non-mutually exclusive mechanisms. The endophyte may fail to colonize a newly produced vegetative tiller, it may fail to colonize the developing seeds prior to dispersal, and it may be lost from successfully colonized seeds prior to germination (Afkhami and Rudgers, 2008, Rudgers et al., 2009). Because we are modelling individual plants, we are unconcerned with the failure of the fungus to colonize some tillers on an otherwise infected plant. We are only concerned with the failure of the fungus to give rise to a new, infected plant. This may happen because the seed produced is not successfully colonized prior to formation and dispersal, or the fungus may die in the seed prior to germination, or the fungal infection may fail to establish itself at the seedling stage prior to developing into a mature plant. We subsume all of these mechanisms into the single parameter *ℓ*. Thus, a positive loss rate means that the proportion *ℓ* of the per capita birth rate from infected individuals will turn out to be uninfected. And 1 – *ℓ* will turn out to have the infection. The special case of *ℓ* = 0 implies that the endophyte is always perfectly transmitted to the next generation. When *ℓ* = 1 the endophyte is strictly horizontally transmitted.

### 2.5. Horizontal transmission

In the case of *strictly* horizontal transmission, we assume that all births from infected plants are initially uninfected (*ℓ* = 1). Furthermore, we assume that the horizontal transmission rate depends on the interactions between infected and uninfected individuals. For any encounter, the chance of passing on the infection is given by *h* (conceptually similar to *β* in standard susceptible–infected (SI) epidemiological models, see e.g. Anderson and May, 1992). Thus the rate of horizontal transmission is given by: *hxy*. A useful extension to our model would be to relax this assumption by creating a spatially explicit model with a more mechanistically detailed treatment of horizontal transmission.

### 2.6. General model

We can use the above to construct a single model of the *Epichloë*–grass interaction as follows:

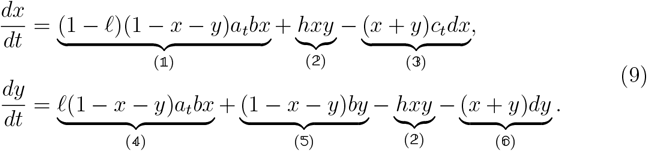

Here, 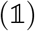 denotes the birth rate at which new infected individuals arise from infected plants, 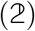 denotes the rate of horizontal transmission by which uninfected plants become infected, 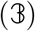 denotes the density-dependent death rate of infected plants, 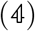 denotes the birth rate at which new uninfected individuals arise from infected plants, 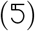 denotes the birth rate at which un-infected individuals arise from uninfected plants, and 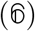 denotes the densitydependent death rate of uninfected individuals. Note that if *ℓ* = 1 we obtain a model that represents strictly horizontal transmission, and if *h* = 0 we obtain a model that represents strictly vertical transmission. Also note that if we let *a_t_* = *a* and *c_t_* = *c*, we obtain a model with temporally constant endophyte (dis)advanatges.

Before analyzing in detail the model results for each mode of transmission, we first examine the dynamics of the system for each of the endophyte × environment interactions shown in Fig. 3.

## 3. System dynamics

Figure 3 can be thought of as depicting an endophyte’s (dis)advantages in six different environments.^4^ In environments Fig. 3a, 3b and 3c the endophyte is a mutualist (i.e. 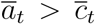), whereas in Fig. 3d, 3e and 3f the endophyte is a parasite (i.e. 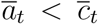). We simulated Eq. (9) for 100 years, for each endophyte-environment combination, for both constant (dis)advantages and for temporally varying (dis)advantages. In Fig. 4 we show the phase portraits for each mode of transmission. From the figure we see that for constant *a* and *c* the system yields point equilibria, while for temporally varying *a_t_* and *c_t_* the system generally yields cyclic equilibria that oscillate around the corresponding point equilibria. We can also see that if the endophyte is strictly vertically transmitted but also a parasite, it goes locally extinct. Finally, it is worth noting the general similarity in the remaining solutions. They all cycle through time. The similarity in the dynamics between the modes of transmission suggests that, in an actual grassland, little can be inferred about the nature of the *Epichloë*-grass relationship directly from an examination of endophyte frequency through time (Rudgers et al., 2009).

**Figure 4:**
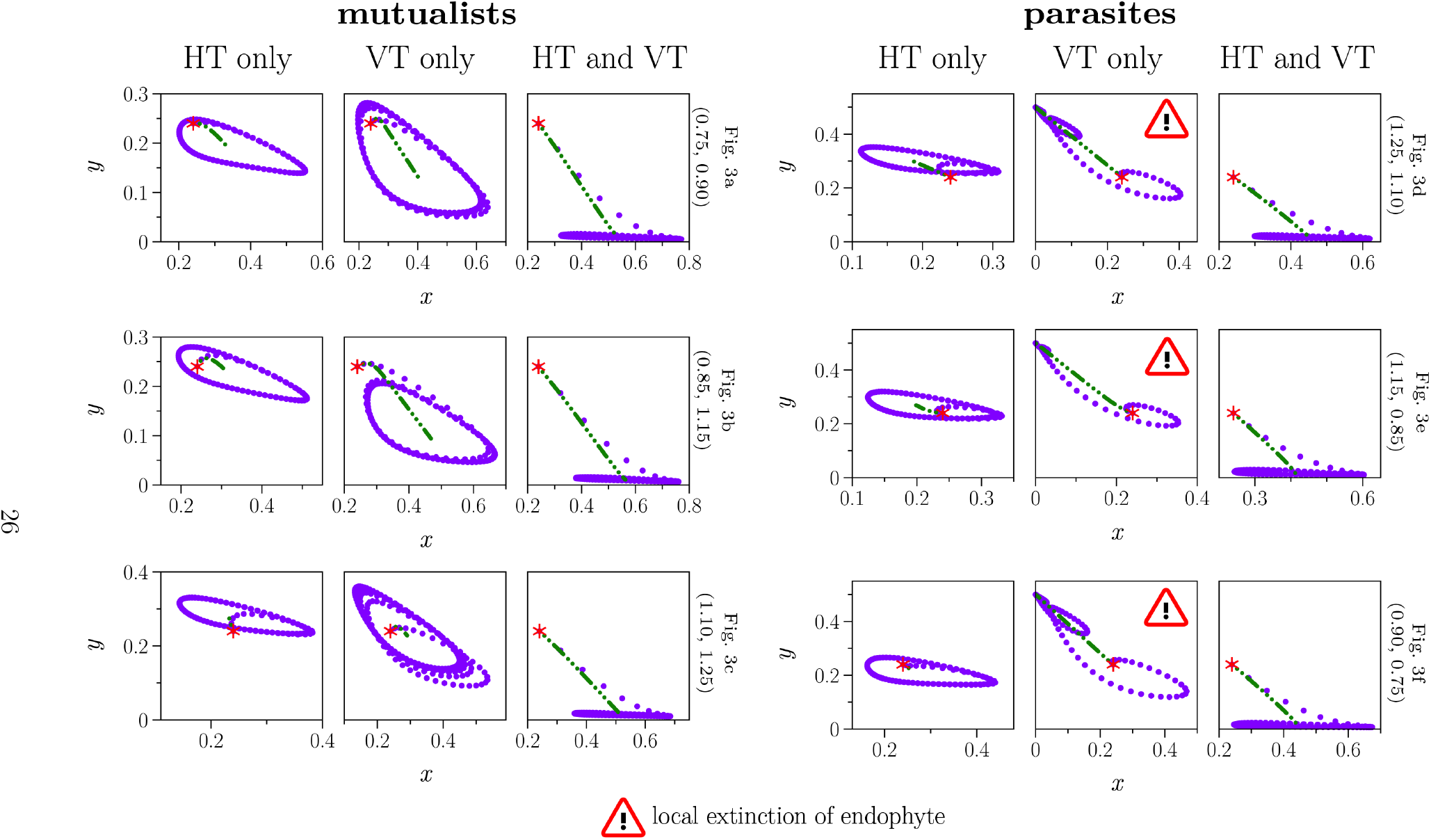
Phase plane diagrams of the system dynamics. HT: strictly horizontal transmission (Case A); VT: strictly vertical transmission (Case B); HT and VT (Case C): both horizontal and vertical transmission. Each set of three corresponds to an endophyte depicted in Fig. 3. The (#,#) denote that (*β*_4_, *α*_4_) values used for the endophyte. See also Fig. 2. The purple dots represent the dynamics for temporally varying 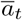 and 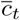. The green dashed-dotted line denotes the corresponding dynamics when 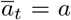 and 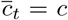 are constants. The panels denoted with the ! symbol indicate that the endophyte goes locally extinct (all are strictly vertically transmitted endophytes that confer a net disadvantage—i.e., they are parasites). The red * in each panel denote the initial conditions.

Next, we use this model to develop each of the three transmission cases separately. For each of these cases we first develop the results for the temporally constant birth and death (dis)advantages, and then we numerically solve for the temporally varying conditions.

## 4. Case A. Strictly Horizontal Transmission, *ℓ* = 1

### 4.1. Case A. Constant (dis)advantages, a_t_ = a, c_t_ = c

In this first case we consider the general situation for the species: *Epichloë baconii, E. glyceriae*, or *E. typhina* sub. *clarkii* (see Table 1). These species seem only to use horizontal transmission; vertical transmission seems to be absent. We obtain this model by setting *ℓ* = 1 in Eq. (9). The following biological constraints exist: 0 < *a*, 0 < *b*, 0 < *c*, 0 < *d*. Subject to these constraints, this model yields two solutions:

#### 4.1.1. Solution A.1. The pasture is totally uninfected at equilibrium

This solution occurs in the case of *h/d* ≤ *c*. In this case the infection cannot sustain itself because infected plants die faster than the infection can be passed on to susceptible plants. The equilibrium pasture densities are given by Eq. (10). Again, note that *x* + *y* need not equal unity.

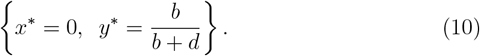

#### 4.1.2. Solution A.2. Infected and uninfected plants coexist at equilibrium

When *c* < *h/d*, the equilibrium proportions of infected and uninfected plants in the pasture are given by Eq. (11).

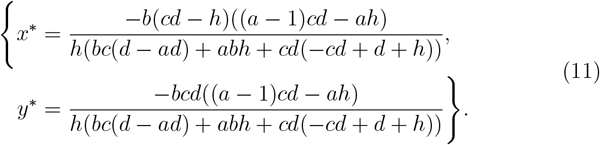

Substituting Eq. (11) into Eq. (1) and simplifying yields:

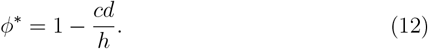

Note that this solution is independent of any birth rate (dis)advantage. Since there is no vertical transmission, any effect the endophyte has on the host plant’s birth rate only influences the birth rate of uninfected plants. Equation 12 is shown in Fig. 5. For now, just concentrate on the lines and ignore the shading surrounding the lines. We see that, *ceteris paribus*, the larger the value of horizontal transmission rate (*h*) or the smaller the value of the death rate (dis)advantage (*c*) the larger the equilibrium fraction of the pasture that is infected (*ϕ*\*). This matches our intuition. Note that we held *a* = 0.9 constant for the results shown for constant *a* and *c* in Fig. 5. This means that, in these examples, the endophyte is always a disadvantage in terms of the host plant’s birth rate. For the area shaded in grey in the figure, the endophyte is also a disadvantage in terms of the host plant’s death rate. Therefore solutions in the grey shading represents endophytes that are parasites. When *c* < 1 the endophyte confers an advantage in terms of the host plant’s death rate. In the yellow shaded region this death rate advantage outweighs the birth rate disadvantage and the endophyte is a mutualist. In the region 0.9 ≤ *c* ≤ 1 (shaded in cross hatching) the endophyte confers a death rate advantage, but that advantage does not outweigh the birth rate disadvantage and so the endophyte is still a net disadvantage and hence a parasite.

**Figure 5:**
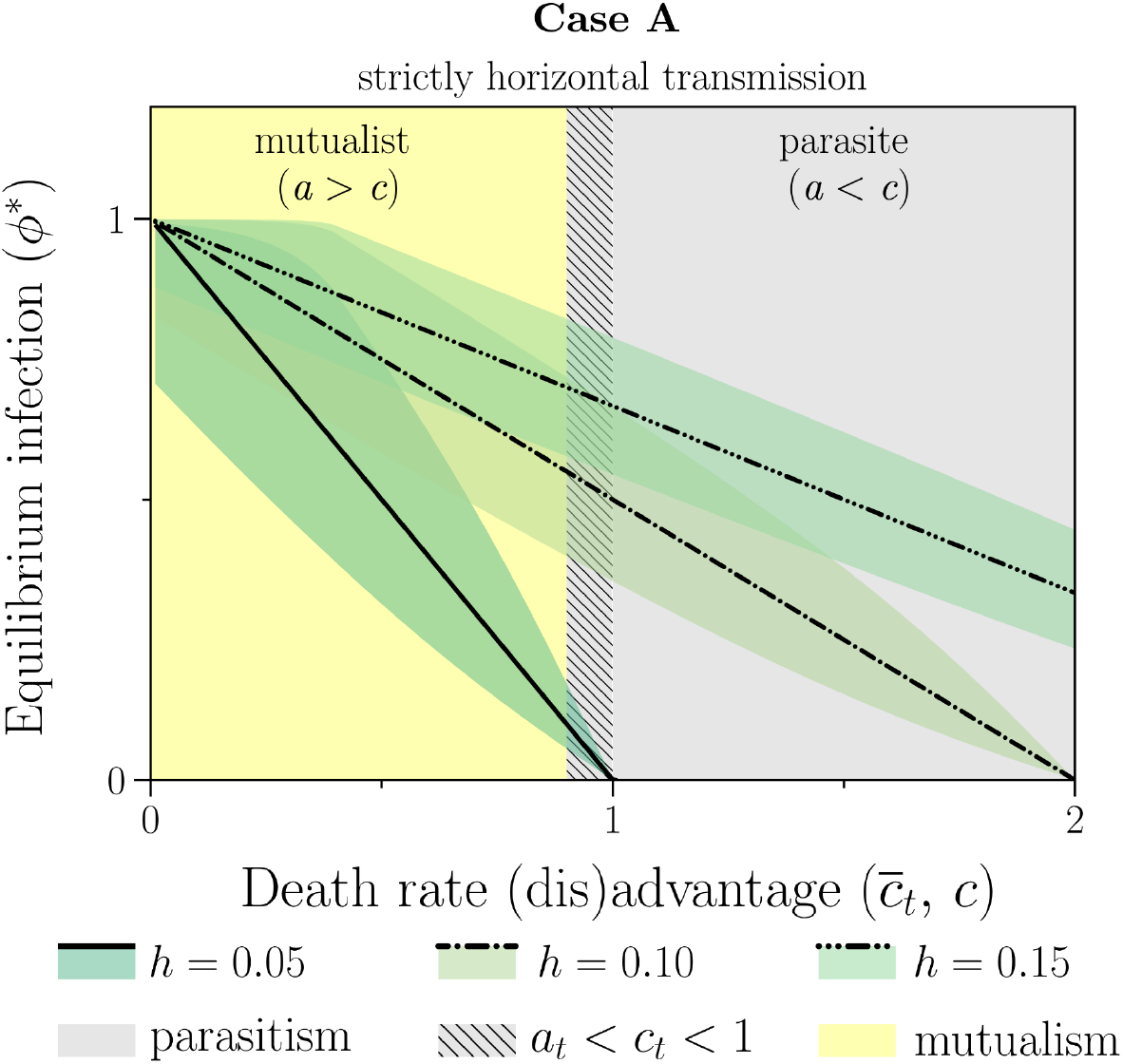
Equilibrium proportion of infected plants, *ϕ*\*. **Case A** (strictly horizontal transmission, *ℓ* = 1), see Eq. (12). The equilibrium proportion of endophyte infected individuals in the population as a function of the death rate (dis)advantage (*c*) conferred by the endophyte. Note that the equilibrium is independent of any birth rate (dis)advantage (*a*) the endophyte might confer since there is no vertical transmission in this case. *h* denotes the rate of horizontal transmission. The shading denotes the minimum and maximum values of the cyclic dynamics for Eq. (9) while the lines denote the solutions when 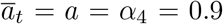 and 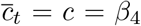 (varied across the *x*-axis) are held constant.

This solution requires that the denominator in Eq. (11) must not be zero. Given the biological constraints on the parameters listed above, this condition is always satisfied. It can also be shown (see Appendix B) that these solutions are all stable fixed points.

### 4.2. Case A. Temporally varying (dis)advantages, a_t_ and c_t_

We simulated Eq. (9) for temporally varying *a_t_* and *c_t_* that correspond to the constant (dis)advantages depicted in Fig. 5. We simulated 100 years of population dynamics, which was sufficient to flush the initial conditions and for the system to settle into its equilibrium dynamics. These results are shown in Fig. 5 as the green shaded regions representing the minimum and maximum values of the cyclic dynamics. Examples of these cyclic dynamics were shown in Fig. 4. From these figures we see that, although the equilibrium dynamics are cyclic (due to the temporally varying *a_t_* and *c_t_*) the cycles oscillate around the equilibrium from the stable point solutions found in the case where *a_t_* = *a* and *c_t_* = *c* are constants. We can see this more generally in Fig. 5, where the point equilibria for the constant (dis)advantage case overlay the stable cycles for the temporally varying *a_t_* and *c_t_*.

## 5. Case B. Strictly Vertical Transmission, *h* = 0

In this section we consider the case for those *Epichloë* species that are strictly vertically transmitted (see Table 1). In comparison to the previous case, we remove the horizontal transmission rate (*h* = 0) and we now consider that infected plants can give rise to infected plants via the vertical transmission of the fungus into the developing seeds (0 ≤ *ℓ* < 1).

### 5.1. Case B. Constant (dis)advantages, a_t_ = a, c_t_ = c

We first analyze this case for perfect endophyte transmission (*ℓ* = 0) and then we consider the more realistic situation of imperfect transmission.

#### 5.1.1. Case B.i. perfect transmission, ℓ = 0

If *ℓ* = 0 it can be shown that the following three solutions exist.

*Solution B.i.1. The pasture is totally infected at equilibrium*. For *c* < *a* the endophyte infection goes to fixation:

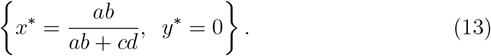

*Solution B.i.2. The pasture is totally uninfected at equilibrium*. For *a* < *c* the endophyte infection goes locally extinct:

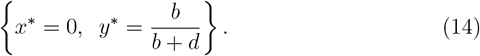

*Solution B.i.3. Infected and uninfected plants coexist at equilibrium*. If, and only if, the per capita birth rate (dis)advantage equals the per capita death rate (dis)advantage, the system yields a coexistence solution—one where both infected and uninfected plants stably coexist in the population. That is, when *a* = *c* we obtain:

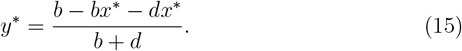

#### 5.1.2. Case B.ii. imperfect endophyte transmission, 0 < ℓ

In the case of imperfect endophyte transmission (i.e. 0 < *ℓ*) the system has two just solutions, either the endophyte goes locally extinct or infected and uninfected plants coexist. No matter how advantageous the endophyte is, there is no solution where the pasture is totally infected [see Eq. (13)] as long as there is imperfect endophyte transmission.

*Solution B.ii.1. The pasture is totally uninfected at equilibrium*. When 0 < *ℓ* and *a* < *c* (i.e. below the diagonal line in Fig. 2) the endophyte infection cannot be maintained in the population. We thus obtain the following equilibrium:

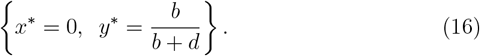

This result demonstrates that when the costs of the endophyte to the host > plant’s fitness outweigh the benefits, the endophyte goes locally extinct. I Note, this is the same equilibrium as found in Eq. (14) above.

*Solution B.ii.2. Infected and uninfected plants coexist at equilibrium*. When 0 < *ℓ* < (*a* – *c*)/*a* (which requires that *c* < *a*) infected and uninfected host plants coexist according to:

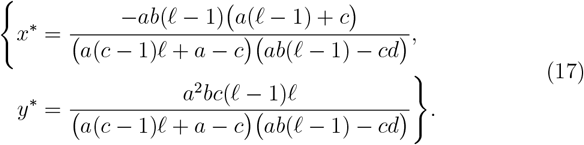

Equation 17 requires that *c*(*b* + *d*) ≠ 0 and *bd*(*a*(*c* – 1)*ℓ* + *a* – *c*)(*ab*(*ℓ* – 1) – *cd*) ≠ 0. Given the constraints on the parameters, it can be shown that these conditions are always met (see Appendix C). Substituting Eq. (17) into Eq. (1) and simplifying yields:

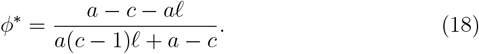

Equation 18 is plotted in Fig. 6a and 6d. There, we see that *ϕ*\* → 0 as *c* → *a* – *aℓ* reflecting the constraints that *ℓ* < (*a* – *c*)/*a*. Notice that there are no solutions where both *a* < 1 and 1 < *c* simultaneously (Fig. 6a grey shaded area). The larger the value of *a*, the larger can be *c* and still result in *ϕ*\* > 0 (*cf*. Fig. 6a and 6d).

**Figure 6:**
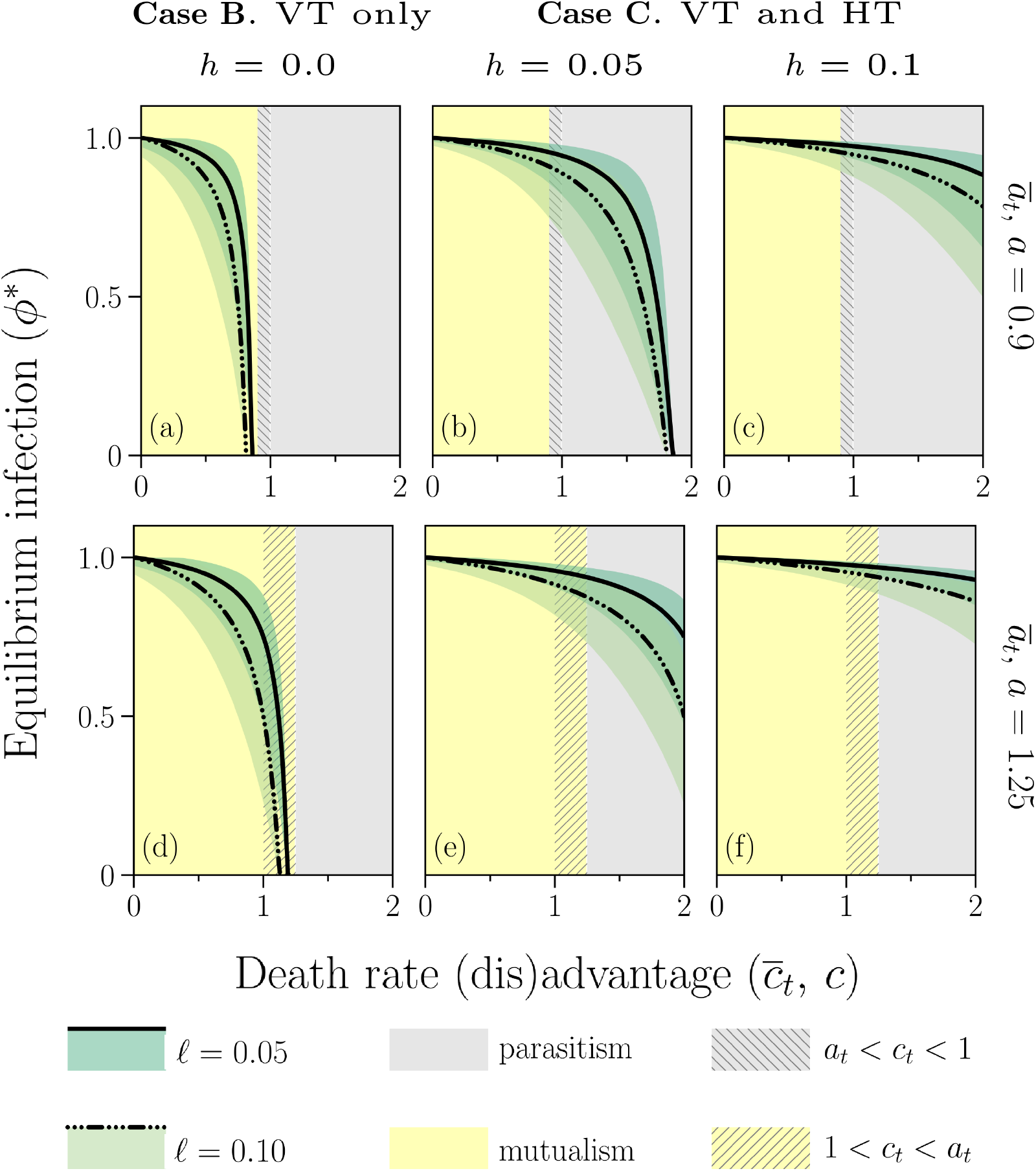
Equilibrium proportion of infected plants, *ϕ*\*. **Case B** (strictly vertical transmission, *h* = 0) is depicted in: (a) and (d). **Case C** (horizontal and vertical transmission, *h* > 0) is depicted in: (b), (c), (e), and (f). Temporally varying (dis)advantages are depicted in all panels; these results are represented by the colored shading. The shading represents the maximum and minimum values between which *ϕ*\* fluctuates at equilibrium (see also Fig. 4). See Eq. (6) for definitions of *α*_1_ and *α*_4_ and these are illustrated in Fig. 3.

### 5.2. Case B. Temporally varying (dis)advantages, a_t_ and c_t_

We simulated Eq. (9) for patterns of variation in *a_t_* and *c_t_*. Again, we simulated the dynamics for 100 years which was sufficient to flush the initial conditions and for the system to settle into its equilibrium dynamics. These results are shown in Fig. 6a and 6d and examples of the dynamics are shown in Fig. 4. As was the conclusion for Case A, we can see from the figures that, although the equilibrium dynamics are cyclic in the case of coexistence (due to the temporally varying *a_t_* and *c_t_*) the cycles oscillate around the equilibrium from the stable point solutions found in the case where *a_t_* = *a* and *c_t_* = *c* are constants. We see this more generally in Fig. 6a and 6d, where the equilibria point solutions for the constant (dis)advantage case overlay the cycles for the temporally varying case.

## 6. Case C. Horizontal & Vertical Transmission, 0 ≤ *ℓ* < 1, 0 < *h*

In this section we consider the case for those *Epichloë* species that simultaneously exhibit both horizontal and vertical transmission (see Table 1).

### 6.1. Case C. Constant (dis)advantages, a_t_ = a, c_t_ = c

Eq. (9) does have a closed-form solution, but it is not enlightening (it involves dozens of polynomial terms). We therefore provide numerical solutions. These solutions are plotted in the second and third columns of Fig. 6. Numerically, we find that coexistence does not require mutualism. An example of such a parasitic interaction is shown in the grey shaded area (where 1 < *c*) of Fig. 6b and 6c (where *a* = 0.9). There we see that coexistence is not only possible, but that the level of pasture infection can even be quite high. *Ceteris paribus*, the higher the horizontal transmission rate (*h*) the higher the fraction of the pasture that is infected (*ϕ*\*; *cf*. Fig. 6b and 6c, or Fig. 6e and 6f). The higher the endophyte loss rate (*ℓ*), the lower the fraction of the pasture that is infected (compare the solid and dashed-dotted lines in any panel). The lower the value of *a* or the higher the value of *c*, the lower the equilibrium fraction of the pasture that is infected, *ϕ*\*.

It can also be shown numerically that all of the solutions in Fig. 6 are stable fixed points; see Appendix B for more detail.

### 6.2. Case C. Temporally varying (dis)advantages, a_t_ and c_t_

Again, we evaluated Eq. (9) numerically for temporally varying *a_t_* and *c_t_*, for a period of 100 years, which was adequate to flush the initial conditions and for the system to settle into its long-term dynamics. Examples of these dynamics are shown in Fig. 4. More generally, the results are shown in Fig. 6. Note again that the solutions to Eq. (9) are cyclic, and are thus illustrated in Fig. 6 as shaded areas that denote the minimum and maximum value of the oscillations. The solutions to Eq. (9) using *a_t_* and *c_t_* are nearly identical to those found using 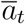 and 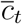, and the same general conclusions hold.

## 7. Discussion

The model clearly demonstrates the following conclusions:

1. For endophytes that are *strictly horizontally transmitted*, the endophytes may be parasites (i.e. in this case, 1 < *c*) and still persist in the population (Fig. 5). Persistence does not *require* the endophyte to be a parasite (i.e. solutions in the left hand side of the graphs where *c* < 1), but a sufficiently high horizontal transmission rate (*h*) *allows* the endophyte to persist while still being a parasite (1 < *c*); see Eq. (12). In this situation it might make sense to talk about a ‘mutualism–parasitism continuum’.
2. For endophytes that are *strictly vertically transmitted*, endophyte per-sistence requires that *c* < *a*. In other words, the advantages of endophyte infection must outweigh the disadvantages of the infection. Put differently, persistence *requires* that the endophyte be a mutualist. Even if we can measure disadvantages of the infection the *net effect* of the endophyte must be an advantage to achieve persistence (*cf*. the grey shaded regions of Fig. 6a). Furthermore, this conclusion still holds when we allow temporal variability in the magnitudes of the advantages and disadvantages, to include periods when there are only disadvantages (*a* < 1 and 1 < *c*, see Fig. 6).
3. For endophytes that are *both horizontally and vertically transmitted*, we reach similar conclusions to those in the strictly horizontal transmission case. Sufficiently large horizontal transmission rates allow the endophyte to persist in the population while being a parasite, although there is no requirement for them to be so (see Fig. 6).

Bear these conclusions in mind as we revisit our considerations of Fig. 3 in the next subsection.

### 7.1. Inferences from ‘experimental’ results

We think that the confusion surrounding the ‘mutualism–parasitism continuum’ is illustrated in the questions we asked earlier about Fig. 3. Here we return to reconsider that figure in light of the model results. In our earlier discussion of this figure we asserted that the endophytes in (a), (b) and (c) are mutualists and that in (d), (e) and (f) they are parasites. We were able to make these assessments based on a quantitative analysis of the (dis)advantages integrated over an entire period of the oscillations (1 year in our examples) for each sine curve. In practice we are unlikely have access to such detailed information. More commonly we would have evidence of some short-term costs and benefits of infection and the previous work of others to draw upon for our inferences. That we are able to measure costs of harboring the endophyte is not particularly surprising; even mutualisms have costs. Nor is it particularly surprising that we might even observe periods of a *net cost* of the endophyte; i.e. a time when the costs outweigh the benefits. What are we to make of such observations? Some might be tempted to label such observations as examples of parasites, or that coupled with previous work that showed benefits, we might be tempted to label the endophyte as part of a mutualism–parasitism continuum. For example, Müller and Krauss (2005, p. 452) state^5^ (references deleted):

> “Stan Faeth and collaborators conclude that most associations between *Neotyphodium* and grass species in natural systems are rarely defensive mutualistic…. Given the strong direct dependence of *Neotyphodium* on its host plant and the only indirect benefits received by the host, a continuum from parasitism to mutualism also appears to be a reasonable description of grass-*Neotyphodium* interactions. The symbioses between *Neotyphodium* and pasture grasses ([tall fescue] and [perennial ryegrass]) in agroenvironments are mostly mutualistic, presumably because the naturally occurring parasitic and potentially more-labile associations have been purged through strong artificial selection.”

What is missing from such inferences is any consideration of the mode of transmission. We argue that such inferences might be justified if the endophyte possesses horizontal transmission (either strictly or in addition to vertical transmission) but it is not justified for strictly vertically transmitted endophytes. To observe a net cost of the endophyte in a strictly vertically transmitted species and infer parasitism or a continuum flies in the face of theory. As Rutherford Aris (1994, p. 127) once said: “Thus the attitude of never believing an experiment until it’s confirmed by theory has as much to be said for it as that which never believes a theory before it is confirmed by experiment.” Theory tells us that over the lifetime of the grass, the endophyte cannot be a net disadvantage and still persist in the environment. It makes sense to speak about a mutualism–parasitism continuum of interspecific interactions between epichoid fungal species and their host grass species when either: (a) we are talking about the genus as a whole, or (b) we are talking about individual *Epichloë* species that have the ability to transmit themselves horizontally. *Pace* Faeth et al. and Müller and Kraus, it does not make sense to describe *Epichloë* species that are strictly vertically transmitted as anything other than mutualists. There may well be times when researchers are able to measure *costs* (or disadvantages as we have called them throughout this paper) to the host plant of this interaction, even at times *net costs*. For such strictly vertically transmitted endophytes, if there exist environments where the net effects of the endophyte are always negative, then the endophyte will not persist in the environment. Some might argue that in such an environment the endophyte is indeed a parasite. We argue that such a classification is both wrong and misleading. At best, one might plausibly claim that in such a situation the endophyte is in an *unstable*, *transient*, or *failed* parasitism.

### 7.2. Model limitations and useful future extensions

All models are simplifications or abstractions. We think our abstraction captures the most important general mechanisms and therefore offers a reasonable approximation to the biological system it is meant to represent. Others might quibble with this assertion. Certainly there are important aspects of the biology that are not incorporated in our model. For example, there is no representation of the seed bank dynamics. The model lacks a stage or age structure. The model also has not been explicitly parameterized for any specific host-endophyte pair, nor for any specific geographical location. By keeping the model non-specific in these ways, we think it demonstrates some useful generalizations. Whether adding any of these biological realisms affects the general conclusions remains to be seen, but our intuition is that the general requirements for coexistence will hold, at least in a qualitative way.

One well known mechanism by which these endophytes may confer an advantage to the host plant is through increased drought tolerance (White et al., 1992). A drought gradient has long been hypothesized to account for the distribution of endophyte-infected perennial ryegrass (*Lolium perenne*) in Europe (Lewis et al., 1997). We will explore this mechanism in future work.

Another useful extension, not previously considered in any rigorous way, would be to explore the role of environmental stochasticity on coexistence. There are plenty of examples in the literature in which stochastic models lead to different, sometimes deeper, understandings than the corresponding deterministic model (see e.g., Newman, 1991, Black and McKane, 2012).

### 7.3. Comparisons with previous modeling work

Ecologist Richard Levins (1966) famously stated:

> “However, even the most flexible models have artificial assumptions. There is always room for doubt as to whether a result depends on the essentials of a model or on the details of the simplifying assumptions….
>
> Therefore, we attempt to treat the same problem with several alternative models each with different simplifications but with a common biological assumption. Then, if these models, despite their different assumptions, lead to similar results we have what we can call a robust theorem which is relatively free of independent lies.” (p. 423)…
>
> “Unlike the theory, models are restricted by technical considerations to a few components at a time, even in systems which are complex. Thus a satisfactory theory is usually a cluster of models. These models are related to each other in several ways: as coordinate alternative models for the same set of phenomena, they jointly produce robust theorems; as complementary models they can cope with different aspects of the same problem and give complementary as well as overlapping results; as hierarchically arranged “nested” models, each provides an interpretation of the sufficient parameters of the next higher level where they are taken as given.” (p. 431)

In this subsection, we review previous modelling attempts for the *Epichloë*-grass interaction, and compare and contrast our models to the previous work. We conclude that, taken together, this set of models jointly produces a robust theorem about this system: that strictly vertically transmitted *Epichloë* endophytes must be mutualists.

Clay (1993) developed a very simple fitness-based model, without endophyte loss, to show that for strictly vertically transmitted fungal endophytes with perfect transmission efficiency, any *net* fitness reduction caused by the endophyte results in the loss of endophyte-infected individuals from the population, echoing earlier work (Ewald, 1987, Fine, 1975). Clay used this model to show that the equilibrium proportion of infected individuals in the pasture is either 0 or 1, and the only means of obtaining a mixed solution was for the benefit of the endophyte to exactly balance the costs, a condition Clay thinks is unlikely to occur. These results are confirmed again by our work, *sensu* Eqs. (13) to (15). The addition that our work contributes here is that by placing our model in a population dynamics framework, we are able to demonstrate how, for this very restricted case (*a* = *c*) it is possible to get other proportions in the mixture apart from the ½. that Clay finds. We show that the particular mixed result achieved depends on the host plant birth and death rates (Eq. (15)).

Ravel et al. (1997) develop a discrete time, stage structured population dynamics model. They divide the endophyte effects on the host plant into effects on birth rate, death rate, and competitive ability. They do not examine cases where the endophyte might be a disadvantage to the host plant in terms of birth and death rates, as we do, although they do briefly admit the possibility that the endophyte might leave the host plant at a competitive disadvantage. The result of this possibility, however, is never explored in their results. In our model, we do not represent competition as distinct from the birth and death rate (dis)advantages. In our model, the competitive advantage of the endophyte, if there is one, arises from how it affects the host plants’ birth and death rates. Finally, Ravel et al. consider a stochastic version of their model where the birth rates, death rates, and competitive advantages are sampled each year from a normal distribution but they provide no details on the mean or variance of these normal distributions, nor do they explore how the mean and variance interact to affect the results. They merely conclude that stochastic effects are only important in small populations, something that has long been known in population ecology. Mostly, Ravel et al. use their model to explore the trade-off between the endophyte loss rate (their *μ*, conceptually equivalent to our *ℓ*) and the birth and death rate advantages of the endophyte. In this paper we are primarily interested in how birth and death rate advantages and disadvantages can trade-off against each other to influence the equilibrium infection rate. For us, 0 < *ℓ* is simply a necessary bit of biological realism, the ommission of which has a fundamental effect on the coexistence outcome.

Gundel et al. (2008) modeled a closed annual grass population using a stage-structured, density-independent, periodic, non-stochastic matrix model. We used a non-stage-structured, density-dependent model of a perennial grass population. Like Gundel et al., we examine environments of deterministic seasonality (see Fig. 3). Also like Gundel et al., we assumed that the grass’ vital rates (*b* and *d*) and transmission efficiency (1 – *ℓ*) were constant through time. Gundel et al. used their model to examine how the endophyte loss rate interacts with the ratio of reproductive rates of the infected and uninfected plants to determine the equilibrium infection frequency. Echoing the earlier work, they conclude that:

> “This condition implies that, in a closed population, the endophyte can only persist in the long term if the infection results in some increase in the host plant fitness…. However, such an increase could be very small….” (Gundel et al., 2008, p. 900)

Making different assumptions, and structuring our model differently, we find the same result, the combined benefits have to outweigh the costs (i.e. there has to be an increase in the host plant’s *net* fitness). However, we show that just because one measures a cost in terms of birth or death rates, this *does not* imply that the endophyte can persist in a population as a parasite. Furthermore, Gundel et al. (2008, p. 902) make the important point that this net fitness benefit of the endophyte can be so small that it would be difficult to measure under field conditions, yet this small difference could result in high infection frequencies in pastures. We find that this conclusion is true, but only for perfect transmission (i.e. *ℓ* = 0). Once the endophyte loss rate is positive, small net positive effects of the endophyte on the host lead to coexistence, but at relatively low frequency of infection in the pasture.

Gundel et al. (2008, p. 897-8) observed that “relatively high infection frequencies have been observed in natural grass populations exhibiting little or no evidence that the fungus confers a reproductive advantage to its host (Saikkonen et al., 1998, Faeth and Hamilton, 2006).” Our model provides two possible explanations for this phenomenon. First, such a solution can occur when the endophyte provides a survivorship advantage instead of a reproductive advantage. Second, if costs and benefits of endophyte infection vary within and between years, it would be possible for the pasture to still maintain a high infection rate even if in some seasons, or across some years, the endophyte provides no benefit, or even a short-term net cost. Notice in Fig. 6 there are examples where *a* < 1, i.e. the endophyte is a disadvantage to the host in terms of reproduction, and yet the endophyte still persists in the population, even possibly at relatively high proportions of the pasture depending on the death rate (dis)advantage (*c*).

Finally, Saikkonen et al. (2002) use meta-population dynamics to understand how the endophyte can be a parasite in some places and still persist on the landscape. They find that this is indeed possible so long as the seed dispersal rates are high enough, and there exist sufficient ‘empty’ patches into which the parasite-infected host plant can disperse. As we stated in the introduction, we do not find the meta-population dynamics explanation convincing. While long-distance dispersal of grass seeds is possible, aided by animals for instance, the vast majority of seed dispersal in these grasses is quite local. To us it seems unlikely that such restricted dispersal would be sufficient to drive the meta-population explanation. Nevertheless, our model cannot be directly compared to this model as we consider closed populations and ignore immigration and emigration.

## Acknowledgments

This research was supported by grants to JAN from the Canadian Natural and Engineering Research Council (NSERC) and the Ontario Ministry of Agriculture, Food, and Rural Affairs (OMAFRA). We thank several anonymous referees for helpful comments on an earlier version of this manuscript.

## Appendix A Systematic Literature Survey of Positions

We conducted a systematic review of the literature to examine the frequency and time trajectory of positions held by authors toward the ‘parasitism-mutualism continuum.’ On 5 November 2019, we executed the following search in Web of Science: `(Epichloë OR Neotyphodium) AND (Parasit* OR Mutuali*)`. This search yielded 399 papers published between 1991 and 2019. We eliminated papers that were about the mutualism between pollinating flies and the sexually reproducing *Epichloë* species, or that mention the fungus only superficially (e.g. only in the reference section). We also excluded papers that were not readily available. The remaining 323 papers were examined by searching the text for `parsiti` and `mutuali` and reading what these authors said specifically about the nature of the plant-fungal interaction. Note that some authors used ‘antagonism’ or ‘pathogenic’ (and their derivatives) to describe a ‘parasitic’ relationship. We searched these terms too.

We classified each paper as taking one of five possible positions: (1) parasitic/parasitism, (2) mutualistic/mutualism, (3) unclear or a continuum, (4) admits the possibility of parasitism, or (5) does not take a position (i.e. makes no statements about mutualism or parasitism). Number 1 was an extremely rare position (5 of 326 papers) and we suspect that the authors were using the word ‘parasitism’ differently than others in this field. To be classified as ‘mutualism’ the authors must have made statements that either the relation *is* a mutualism, or that it is *believed to be, thought to be*, etc. a mutualism. To be classified as ‘unclear or a continuum’ we looked for statements such as:

> “Symbioses between cool-season grasses (Poaceae subfamily Pooideae) and fungi of genus *Epichloë* (including their asexual derivatives in the genus *Neotyphodium*) are very widespread and occur in a broad taxonomic range of this important grass subfamily. These symbioses span the continuum from mutualistic to antagonistic interactions” (Schardl et al., 2008, p. 483)

The above quote clearly refers to the genus *Epichloë* as a whole, and is therefore a correct usage of the continuum concept (i.e. it is certainly correct that various species sit at different locations along the continuum). Here is an example of the implied use of the continuum concept when clearly speaking only of the strictly vertically transmitted species; we suggest this is an inappropriate characterization of the nature of the interaction:

> “The negative effect of asexual endophytes on their hosts may evolve because the reward for the partners and the ability to control the interaction is asymmetrical. The fitness of the fungus depends strongly on the fitness of the plant, but the plant does not always benefit from the presence of the fungus. This, together with the fact that the host–endophyte interaction seems to be controlled by the host, makes it likely that the outcome of the interaction is antagonistic rather than mutualistic. A supposedly mutualistic host–endophyte interaction may appear parasitic after detailed investigation. Here, we show that an apparently beneficial asexual endophyte may have a negative effect on the reproduction of its host.” (Olejniczak and Lembicz, 2007, p. 486; references omitted)

A paper was classified as ‘admitting the possibility’ if it referred to the idea that the interaction might be parasitic without appearing to endorse that position. This category might be viewed as a weaker version of the ‘unclear or continuum’ category. Finally, ‘no position’ was used when the authors did discuss the nature of the endophyte–plant interaction.

Fig. A.1 shows the accumulation of papers taking each of the above positions. As of 2019, of the 286 papers that take a position, 70% state that the relationship is a mutualism, and 30% take the position that the relationship is ‘unclear,’ a ‘continuum,’ or at least it is possible the relation is not a mutualism.

**Figure A.1:**
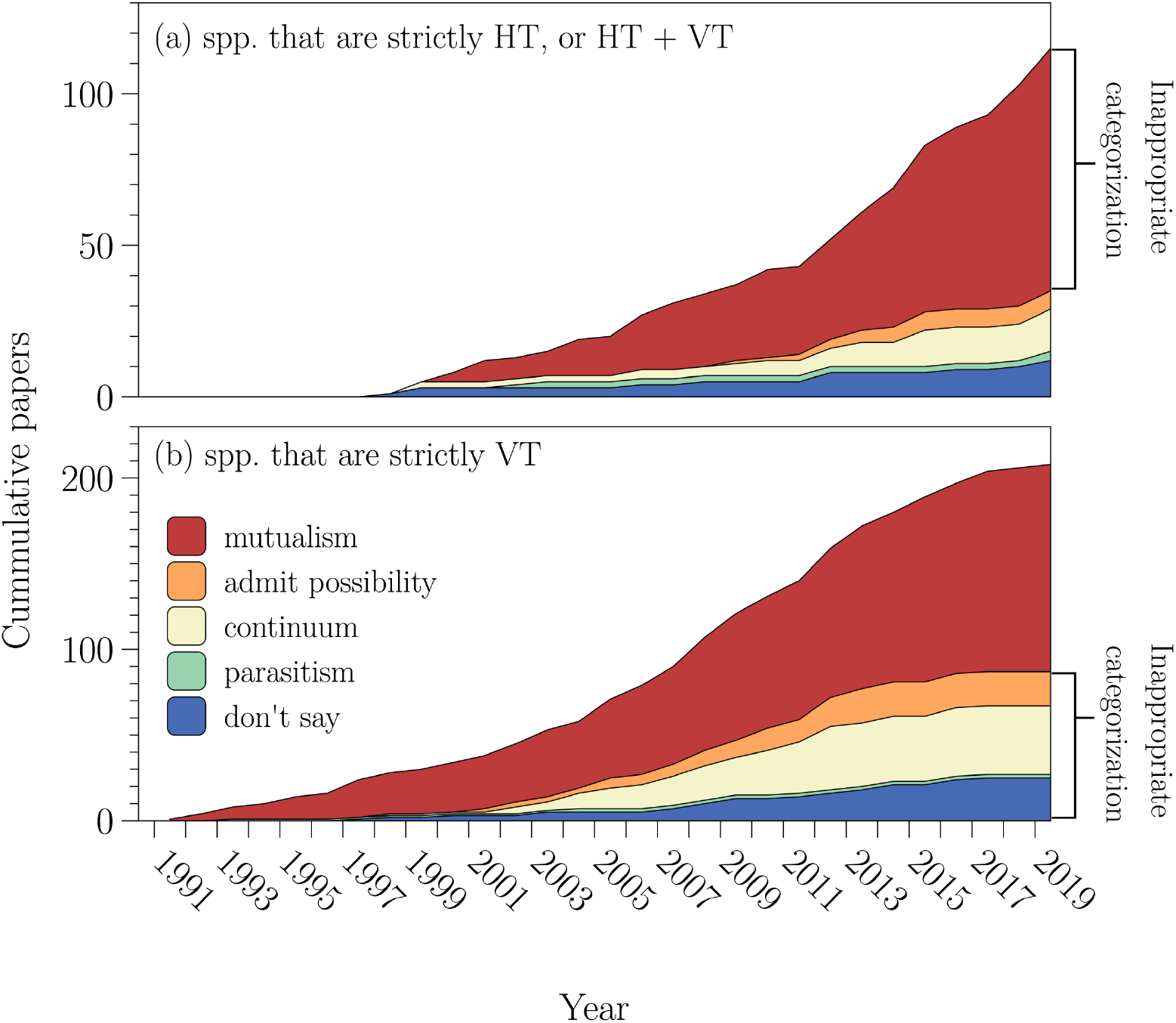
The cumulative number of research papers published on *Epichloid* species, cross classified by the transmission mode(s) of the subject species (according to Leuchtmann et al. 2014), and how the authors describe the relationship between the *Epichloid* species and the host grass plant. (a) denotes the cumulative number of papers that have as their subject matter an *Epichloë* species that is either strictly horizontally transmitted (HT) or has both horizontal and vertical transmission (HT + VT); (b) denotes the same information where the focal species is strictly vertically transmitted (VT). Note that in (a) it is inappropriate to refer to these interactions as mutualisms (they could be, but equally they could be parasites), while in (b) it is inappropriate to refer to these interactions as anything other than a mutualism.

These numbers are confounded with the relative publishing frequency for authors taking the various positions. However, it is not uncommon to see the same authors’ positions shifting through time (e.g. *mutualism*: Newman et al., 2003, Ryan et al., 2014, Shukla et al., 2015, Bastías et al., 2018), (*unclear/continuum/admits possibility*: Hunt and Newman, 2005, Rasmussen et al., 2007, Antunes et al., 2008).

## Appendix B Linear Stability of Fixed Point Solutions

### B.1 General state equations

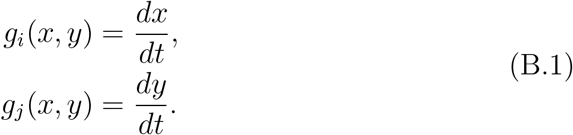

Note that the general equations are equations (6a) and (6b) in the main text, and Cases A and B can be retrieved by simply setting *ℓ* = 1 or *h* = 0 respectively.

### B.2 Linear stability of fixed point solutions

To evaluate the linear stability of the fixed point solutions, we substitute the fixed points into the relevant Jacobian matrix and then calculate the Eigenvalues. If the real parts are both negative then the fixed point is a stable equilibrium. The Jacobian is given by:

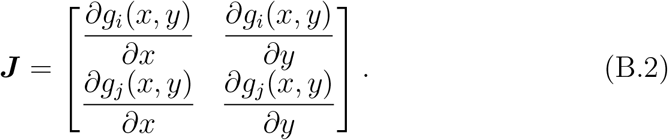

To calculate the Eigenvalues we need:

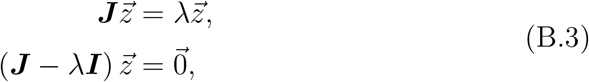

where ***ℓ*** is a 2 × 2 identity matrix and 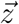 is a vector of length 2. The characteristic equation is then given by:

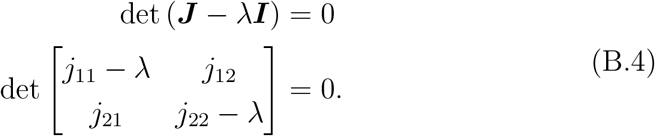

From here we can obtain the characteristic polynomial:

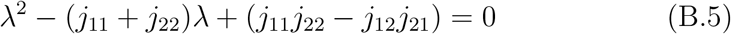

Eq. (B.5) always has two roots. These roots can be real or complex, and they do not have to be distinct. Employing the quadratic formula we can calculate the roots of Eq. (B.5) as:

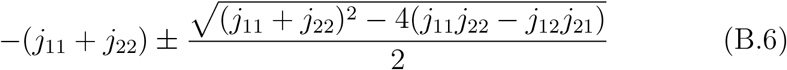

The fixed point solutions to the system of equations are stable if the real parts of the Eigenvalues are both negative.

### B.3 Case A. Strictly horizontal transmission, ℓ = 1

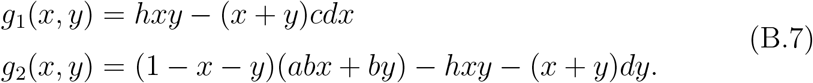

Now calculate the Jacobian matrix. To do this we calculate the partial derivatives:

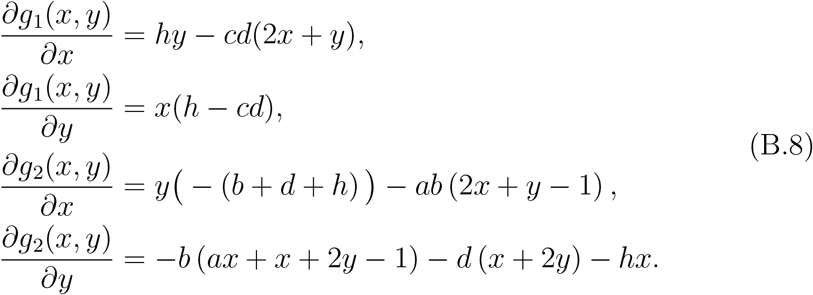

Which yields:

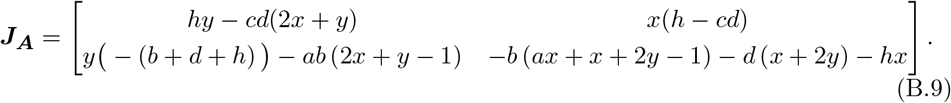

Next we calculate the Eigenvalues:

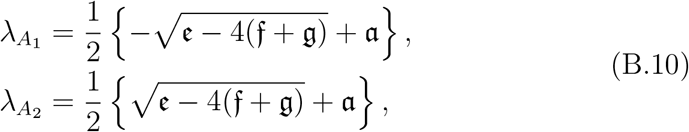

where

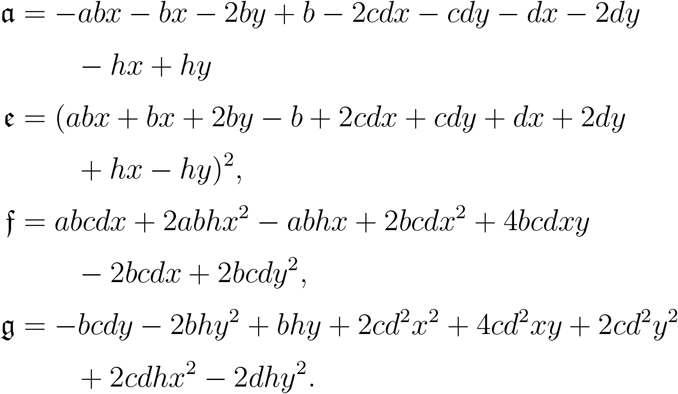

Using the solutions depicted in Fig. 5, it can be easily shown numerically that for these fixed point solutions the real parts of the two Eigenvalues are both negative, and therefore these solutions are stable.

### B.4 Case B. Strictly vertically transmitted, h = 0

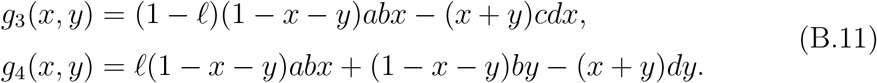

The partial derivatives are then:

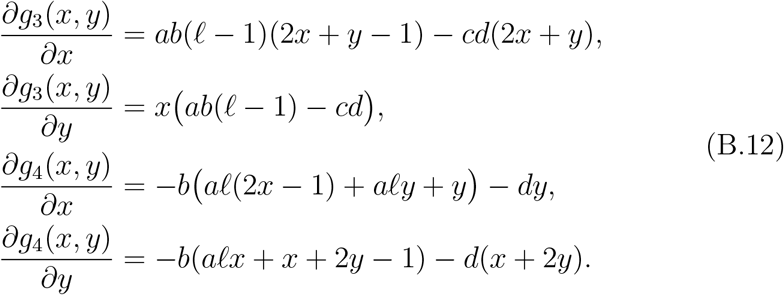

Which yields

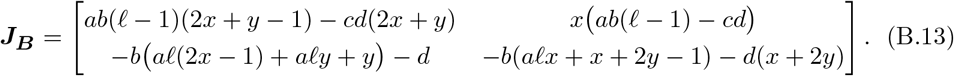

The Eigenvalues are then:

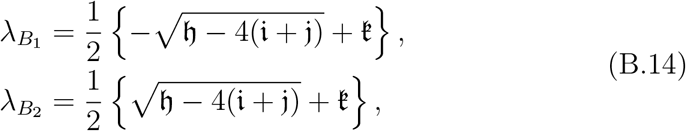

where

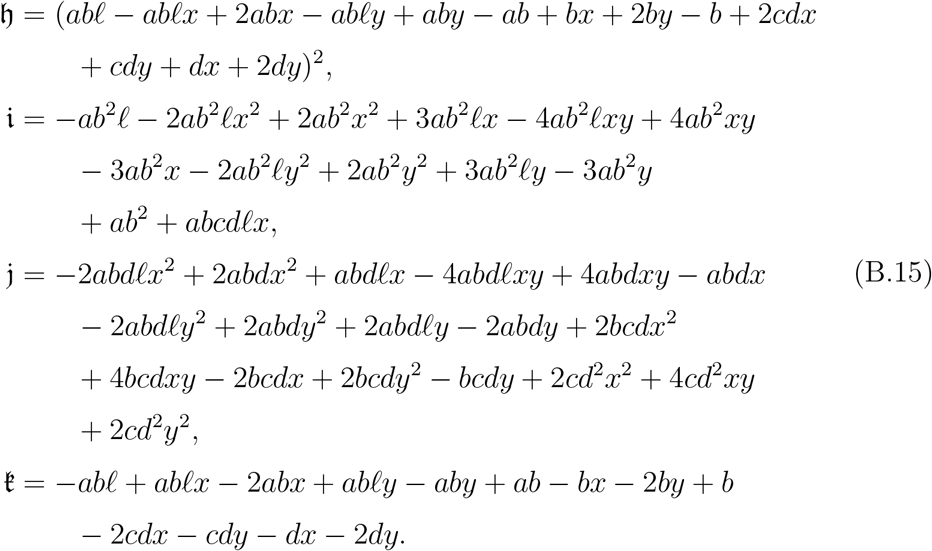

### B.5 Case C. Both horizontally and vertically transmitted, 0 < ℓ < 1 and 0 < h

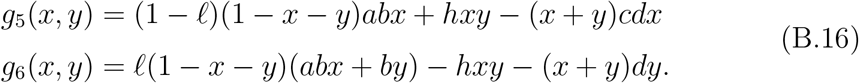

The partial derivatives are then:

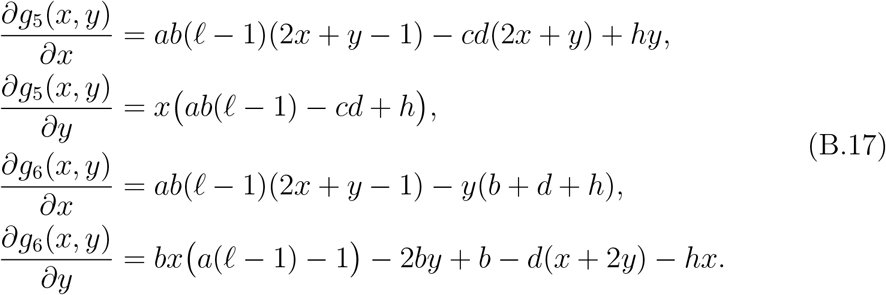

Which yields:

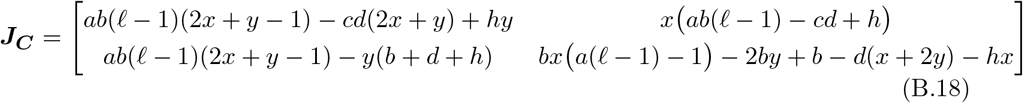

The Eigenvalues are then:

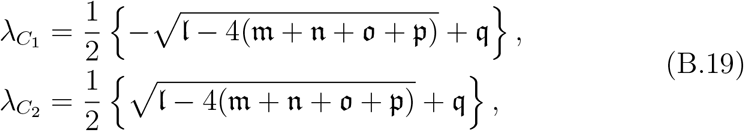

where

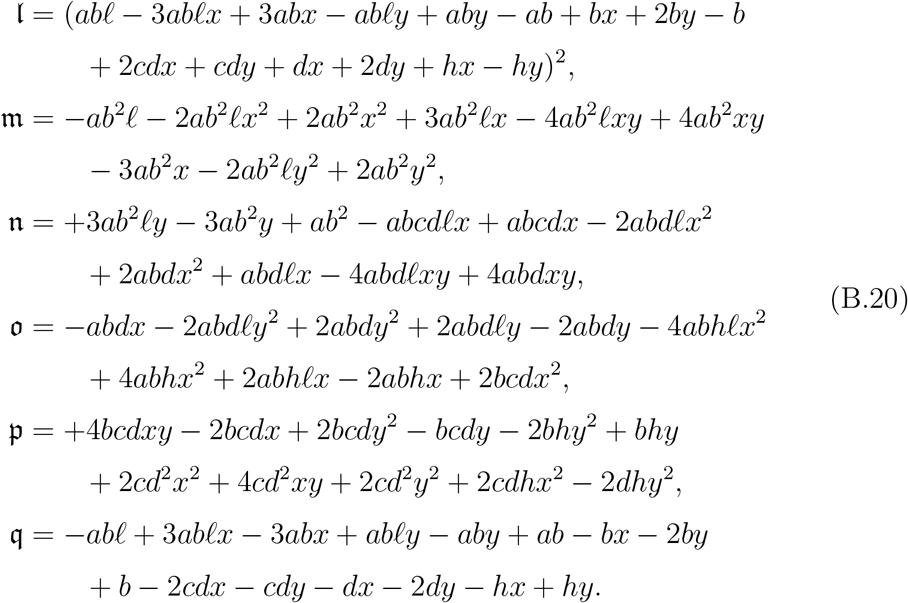

Since we solved Eq. (9) numerically for Case C, we evaluated the stability of these solutions by substituting the numerical solutions into Eq. (B.19) and checking the signs of the real parts of the Eigenvalues. All feasible solutions were stable.

## Appendix C Solution B.ii.2 Conditions

In this appendix, we show that the conditions on the coexistence solution (equation 14) never constrained this solution, given biologically feasible parameter values. Eq. (17) requires that:

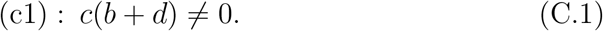

Since 0 < *c*, 0 < *b*, and 0 < *d, c*(*b* + *d*) can never equal 0; so this condition is always met.

Eq. (17) also requires that:

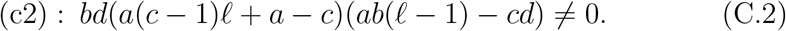

It can be shown that (c2) is met if the following are true:

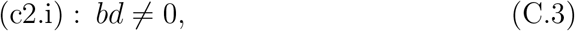

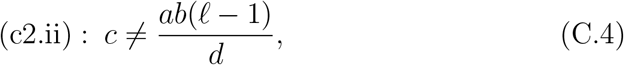

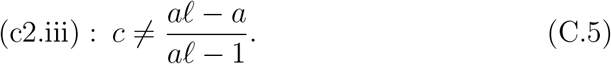

Since 0 < *b* and 0 < *d*, (c2.i) is always true. Since *ℓ* < 1, condition (c2.ii) would require *c* < 0 for it to equal the right hand side of the equation and so, since biologically 0 < *c*, (c2.ii) is always true. Condition (c2.iii) can be *false* in two further sets of conditions:

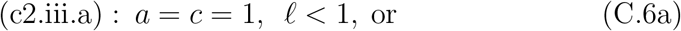

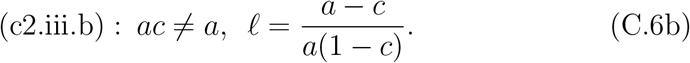

Condition (c2.iii.a) denotes the biologically unlikely case in which the en-dophyte imposes no advantages or disadvantages at all. This case is unlikely since stochastic processes (akin to genetic drift in a population genetics model) would ensure that, eventually, the endophyte would be lost from the system. Next, we will show that the conditions that support a coexistence solution are such that (c2.iii.b) never constrains the solution space, and hence for coexistence, the solution to Eq. (17) can be true subject to the constraints 0 < *ℓ* < (*a* – *c*)/*a* and *c* < *a*.

Using the Eq. (1) for *ϕ*\*, a coexistence solution requires that:

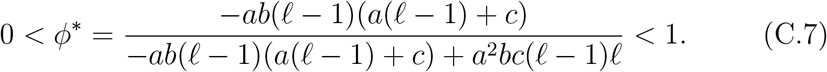

Eq. (C.7) can be simplified to:

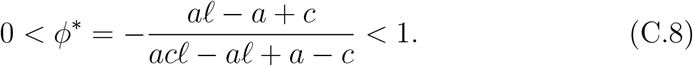

Eq. (C.8) is true iff both of the following condition holds:

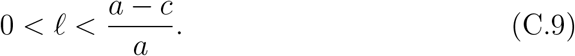

Clearly, a necessary condition for Eq. (C.9) to be true is that *a* > *c*.

## Footnotes

1 Note that all of the asexual *Epichloë* were classified as *Neotyphodium* until 2014. Note as well that, as of the taxonomic revision in 2014, *Neotyphodium starii* is no longer considered an epichloid species (Leuchtmann et al., 2014). It nevertheless is an asexual fungal endophyte that has no apparent horizontal transmission and produces alkaloids in the same classes as the *Epichloë*.

2 Note that mycorrhizal fungi are horizontally transmitted (see e.g. Genkai-Kato and Yamamura, 1999).

3 Note that the phase shift is inconsequential to the qualitative conclusions. *α*_3_ = 0 and *β*_3_ = 600 were chosen for the visiual effect of creating maximal contrast between the temporal patterns of *a_t_* and *c_t_*.

4 Fig. 3 can equally be thought of as representing six different endophyte species in the same environment. The same logic applies and the same conclusions are reached.

5 Recall that Neotyphodium was the genus name for all strictly vertically transmitted *Epichloë* species until the most recent taxonomic revision (Leuchtmann et al., 2014).

